# Telomere-to-telomere assembly by preserving contained reads

**DOI:** 10.1101/2023.11.07.565066

**Authors:** Sudhanva Shyam Kamath, Mehak Bindra, Debnath Pal, Chirag Jain

## Abstract

Automated telomere-to-telomere (T2T) *de novo* assembly of diploid and polyploid genomes remains a formidable task. A string graph is a commonly used assembly graph representation in the overlap-based algorithms. The string graph formulation employs graph simplification heuristics, which drastically reduce the count of vertices and edges. One of these heuristics involves removing the reads contained in longer reads. However, this procedure is not guaranteed to be safe. In practice, it occasionally introduces gaps in the assembly by removing all reads that cover one or more genome intervals. The factors contributing to such gaps remain poorly understood. In this work, we mathematically derived the frequency of observing a gap near a germline and a somatic heterozygous variant locus. Our analysis shows that (i) an assembly gap due to contained read deletion is an order of magnitude more frequent in Oxford Nanopore reads than PacBio HiFi reads due to differences in their read-length distributions, and (ii) this frequency decreases with an increase in the sequencing depth. Drawing cues from these observations, we addressed the weakness of the string graph formulation by developing the RAFT assembly algorithm. RAFT addresses the issue of contained reads by fragmenting reads and producing a more uniform readlength distribution. The algorithm retains spanned repeats in the reads during the fragmentation. We empirically demonstrate that RAFT significantly reduces the number of gaps using simulated datasets. Using real Oxford Nanopore and PacBio HiFi datasets of the HG002 human genome, we achieved a twofold increase in the contig NG50 and the number of haplotype-resolved T2T contigs compared to Hifiasm.

## 1 Introduction

Building high-quality haplotype-resolved *de novo* assemblies remains a principal challenge in genomics research. The T2T (telomere-to-telomere) assembly of CHM13 human genome [23] is a recent scientific milestone which has inspired further efforts toward achieving T2T assembly of personal diploid human genomes [14, 33]. Third-generation read sequencing technologies like Pacific Biosciences high-fidelity (PacBio HiFi) reads and Oxford Nanopore Technology (ONT) reads were instrumental in constructing the CHM13 reference genome. Currently, PacBio HiFi and ONT Duplex sequencing technologies produce reads that have an average length greater than 10 kbp and per-base error rates less than 0.5% [19, 31].

*De novo* genome assembly using long reads is most commonly solved using overlap-layout-consensus based methods [22]. The assembly workflow typically involves (i) computing pairwise overlaps between reads, (ii) error-correction of reads, (iii) constructing a read-overlap graph, and (iv) identifying walks in the graph which correspond to contiguous substrings of the genome. In a read-overlap graph, the reads are represented as vertices and the suffix-prefix overlaps between the reads are represented as directed edges. The initial version of this graph is quite tangled and requires additional graph simplification heuristics to remove redundant vertices and edges. Myers’s *string graph* formulation [21, 22] has long been the standard choice to build a simplified version of a read-overlap graph [2, 4, 5, 6, 7, 8, 9, 15, 17, 28, 29]. The string graph model was also used by the T2T Consortium to assemble the CHM13 human genome [23].

The two important graph-simplification steps in the string graph formulation are (i) removal of transitively inferable edges and (ii) deleting those reads that are entirely contained as substring in another read [21]. The advantage of these two steps is that they prohibit redundancy, i.e., no two walks in a string graph spell the same sequence [27]. However, prior works have highlighted that the removal of contained reads from the graph is an ‘unsafe’ operation because this heuristic can occasionally disconnect the walks corresponding to true chromosome sequences [4, 9, 12, 13, 19, 23]. The connectivity breaks when all reads that cover one or more genomic intervals are removed. We refer to these events as assembly gaps due to contained read deletion (formally defined in Methods). The need to remove contained reads is currently a major weakness of the read-overlap-based assembly algorithms [19].

A few approaches have been proposed to tackle the above problem. The algorithms in [12] and [13] work under simplified assumptions on input read lengths and sequencing coverage. These algorithms don’t trivially extend to practical solutions for assembling highly repetitive genomes. An initial release of the Hifiasm assembler [5] included a method to recover an essential contained read if the read connects the ends of two walks in the graph. This technique has been observed to work in simple scenarios but is not always reliable [13, 19]. A more recent version of Hifiasm also uses alignments of ultra-long nanopore reads to a string graph to identify the necessary contained reads [4]. This is a useful approach if ultra-long reads are available. Previous experiments [13] have reported that contained read deletion in a string graph is more likely to impact graph connectivity in the regions of low heterozygosity. In such regions, a longer read sampled from one haplotype may contain all reads that cover the homologous region in the opposite haplotype.

A sequencing run results in a multiset of reads. Considering all possible distinct sequencing outputs, we mathematically derived a formula to calculate the fraction of sequencing outputs in which an assembly gap occurs due to contained read deletion. It is useful to compare the fractions in different experimental settings, e.g., with different choices of sequencing technology and sequencing depth. We performed this theoretical analysis for both normal and cancer genomes. The analysis reveals novel insights into the key factors contributing to assembly gaps due to contained read deletion. We refer to this method as CGProb (https://github.com/at-cg/CGProb).

Next, using insights from CGProb, we developed RAFT (Repeat-Aware Fragmenting Tool, https://github.com/at-cg/RAFT) to prevent assembly gaps during genome assembly. Conceivably, the proportion of contained reads in a sequencing dataset is roughly determined by its read-length distribution. On the one hand, there are no contained reads if all reads have a fixed length, whereas an ONT sequencing dataset may have a significant fraction of contained reads due to a wide read-length distribution [20]. RAFT reduces the range of read lengths in a sequencing dataset by fragmenting long reads into equal-sized shorter reads. The reads predicted to span repetitive regions of the genome are treated differently. RAFT enables high-quality phased assemblies of variable-length long and accurate reads (e.g., ONT Duplex reads or a mixture of ONT Duplex and PacBio HiFi reads). Both of the above tools, RAFT and CGProb, are useful in the era of telomere-to-telomere genomics.

## 2 Results

### 2.1 Overview of CGProb

Haplotype-resolved assembly of diploid genomes is challenging because one needs to distinguish between reads originating from two near-identical haplotype sequences. The differences between the haplotypes occur at heterozygous loci. Contained read deletion may break haplotype walks in a read-overlap graph. We show an example in Figure 1A where an assembly gap occurs in the second haplotype after the deletion of contained read *r*_8_. For brevity, we refer to the assembly gaps due to contained read deletion as just ‘assembly gaps’ in this section. The occurrence of assembly gaps in a string graph depends on several factors, including the sampling positions of reads, genome heterozygosity, sequencing coverage, ploidy, etc. As a result, deriving the expected number of assembly gaps in a string graph is challenging. Knowing this value for different choices of sequencing technology and sequencing depth can allow a more informed decision-making for *de novo* genome sequencing.

**Figure 1:**
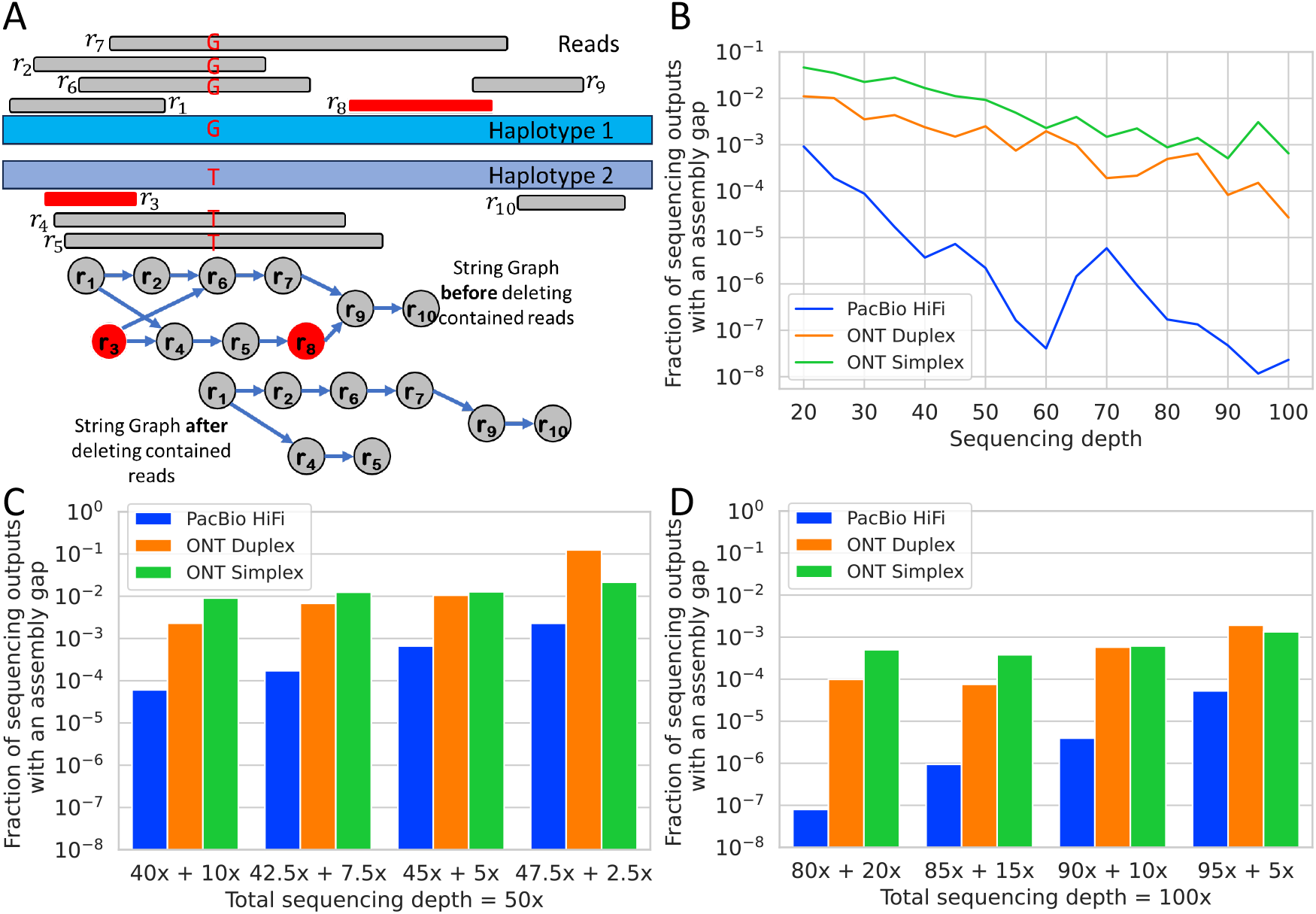
Assembly gaps and their occurrence frequency. **(A)** An example of a sequencing output where an assembly gap occurs in the string graph due to contained read deletion. Read *r*_3_ is contained in read *r*_1_. Read *r*_8_ is contained in read *r*_7_. Accordingly, the string graph representation excludes reads *r*_3_ and *r*_8_. Read *r*_3_ is redundant; its deletion simplifies the graph. However, removing read *r*_8_ breaks the connectivity between reads *r*_5_ and *r*_9_, which was necessary to spell the second haplotype. **(B)** Fraction of sequencing outputs containing an assembly gap. We measured the fractions using the read-length distributions corresponding to three sequencing technologies (PacBio HiFi, ONT Duplex, ONT Simplex) and using different sequencing depths. Here we used equal sequencing depths on both haplotypes. **(C)** and **(D)** Fraction of sequencing outputs containing an assembly gap when the sequencing depths across the two haplotypes are uneven. This scenario models somatic mutation in DNA with variant allele frequency below 0.5. In panel (C), the total sequencing depth for both haplotypes is 50×. In panel (D), the total sequencing depth is 100×.

Consider the output of a sequencing experiment as a multiset of reads, where each read is identified by its haplotype of origin, length, and stop position. The user provides a read-length distribution and sequencing depths of two haplotypes as input. Accordingly, the set of valid sequencing outputs includes all possible multisets of read-sampling intervals that are consistent with the user-provided input (Methods). CGProb considers all these valid sequencing outputs and calculates the fraction of sequencing outputs in which an assembly gap occurs (Methods). We made a few simplifying assumptions to make the analysis feasible: e.g., (i) there is a single heterozygous SNP locus in the diploid genome, (ii) reads are error-free, and (iii) the two haplotypes do not have repetitive DNA (Methods). Although these assumptions will likely not hold in practice, the above model is informative to study the frequency of an assembly gap near an isolated heterozygous locus while assembling error-corrected long reads. A naive method to calculate the fraction would check all possible *O*(*G*^*N*^ ) read sequencing outputs individually, where *G* is the genome length and *N* is the number of reads. Instead, we developed an efficient combinatorial technique to count the sequencing outputs containing an assembly gap in polynomial time. The theory and implementation details are provided in the Methods.

### 2.2 Frequency of observing an assembly gap

Using CGProb, we evaluated the frequency of observing an assembly gap near a heterozygous SNP. In the first scenario, the heterozygous SNP is a germline mutation. Here we used equal sequencing depths for both haplotypes (paternal and maternal). In the second scenario, evidence for a heterozygous SNP is observed in the sequencing output due to a somatic mutation with variant allele frequency^1^ below 0.5. Here, we used uneven sequencing depths for the two haplotypes (e.g., tumour and normal). We considered three prominent sequencing technologies: PacBio HiFi, ONT Simplex, and ONT Duplex. For each sequencing depth and for each technology, we simulated five read length distributions consistent with that sequencing technology and depth of coverage (Methods). We ran CGProb on each read length distribution and recorded the median fraction. The minimum and maximum values are shown separately in Supplementary Figures S1, S2.

#### (1) Germline heterozygous variant locus

We computed the fraction of the sequencing outputs containing an assembly gap while varying the genome sequencing depths from 20×to 100 × (Figure 1B). Here the sequencing depths were balanced equally on both haplotypes, e.g., 20 × depth corresponds to 10 × depth on each haplotype. Our results show that there is at least an order of magnitude difference in the fraction of sequencing outputs containing an assembly gap for ONT reads compared to PacBio HiFi reads. The results imply that assembly gaps are more frequent when there is a larger variation in read lengths. The read-length distribution of ONT reads is generally more skewed than PacBio HiFi reads (Supplementary Figure S3). Intuitively, the fraction of contained reads will be greater if the variation in read lengths is larger.

Figure 1B also shows a decrease in the median fraction as sequencing depth increases, although some deviation from this trend is observed for PacBio HiFi reads due to noise arising from our use of a small number of trials. This decreasing trend is also intuitive because if the sequencing coverage is higher, then the number of times a genome interval is sequenced becomes larger which reduces the chance of every read which supports that interval being a contained read.

#### (2) Somatic heterozygous variant locus

We analysed the fraction of the sequencing outputs containing an assembly gap in a simulated heterogeneous sequencing sample, e.g., a sample with mixed normal and tumour cells. We set total sequencing depths as 50 × and 100 × . We set the tumour sequencing depths as 5%, 10%, 15%, and 20% of the total (Figures 1C, 1D). For all three sequencing technologies, we observed that the fraction of sequencing outputs containing an assembly gap increases as the tumour sequencing depth decreases. The result implies that assembly gaps are more frequent near somatic genetic variants with lower variant allele frequencies. This is expected because all the reads sampled from an interval are more likely to be contained in a read from the second homologous interval if the coverage over the first interval is low and the coverage over the second interval is high. We again found that a string graph of ONT reads is more likely to contain an assembly gap than PacBio HiFi reads.

### 2.3 Overview of RAFT

The above analysis indicates that the problem of assembly gaps due to contained read deletion is much less prevalent with narrow read-length distributions. Inspired by these results, we developed RAFT as a practical solution to assemble a long-read dataset when there is significant variability in the read lengths. The RAFT algorithm fragments long reads into shorter, uniform-length reads while also considering the potential usefulness of the longer reads in assembling complex repeats. We envision RAFT as a module that can be easily integrated into any existing overlap-layout-consensus-based assembler.

The input to the RAFT algorithm includes error-corrected long reads and all-to-all pairwise alignment information (Figure 2A). The algorithm carefully fragments the input reads. While fragmenting a read *r*, we consider its high-quality pairwise alignments with other reads. If the number of alignments to an interval in read *r* exceeds *μ · cov*, where *μ* is a user-specified threshold (default value = 1.5) and *cov* is the coverage of the input sequencing dataset, then we prevent the interval from being fragmented. Such intervals are potentially repetitive and may be necessary to resolve repetitive sequences (Figure 2B). We set the default length of fragmented reads as 20 kbp to ensure that the fragmented read lengths are greater than the lengths of the abundant interspersed repeats such as LINEs (Methods).

**Figure 2:**
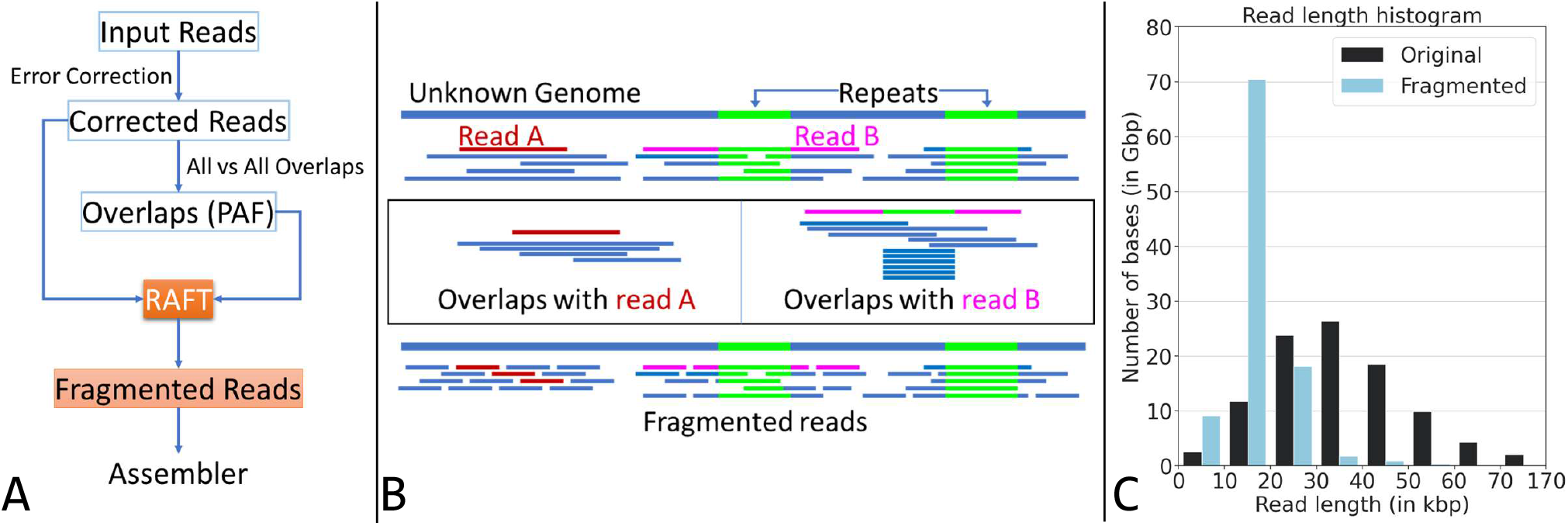
Illustration of the RAFT algorithm and its usage for genome assembly. **(A)** Flowchart of an assembly workflow that uses RAFT. RAFT accepts error-corrected long reads and all-to-all alignment information as input. It produces a revised set of fragmented reads with a narrow read-length distribution. **(B)** Illustration of the RAFT algorithm. Read A (shown in red) is sampled from a non-repetitive region of the genome. Accordingly, RAFT fragments read A into shorter uniform-length reads. Read B (shown in pink) spans a repetitive region of the genome. RAFT detects the repetitive interval in read B because more than the expected number of sequences align to that interval. The portions of read B outside the repetitive interval are split into shorter reads. **(C)** The impact of RAFT can be seen on a set of ONT Duplex reads sampled from the HG002 human genome. The range of the read-lengths is significantly reduced by using RAFT. The original dataset comprises 3.7 million reads with a skewed read length distribution. After fragmentation, the dataset comprises 6.8 million reads.

RAFT fits conveniently in a *de novo* genome assembly workflow in between the long-read error correction step and the assembly steps (Figure 2A). We designed the ‘RAFT-Hifiasm’ workflow that combines RAFT’s ability to manipulate the read-length distribution and Hifiasm’s highly efficient all-to-all alignment and error-correction algorithms [5]. Accordingly, the RAFT-Hifiasm workflow uses Hifiasm for error-correction of input reads and computing all-vs-all pairwise read alignments. RAFT uses this information to generate a set of fragmented reads (Figure 2C). In the end, we assemble the fragmented reads using Hifiasm.

### 2.4 Evaluation using simulated data

We simulated error-free long reads from a publicly available haplotype-resolved HG002 diploid human genome assembly using Seqrequester [30]. We simulated one PacBio HiFi (30 × ), two ONT Simplex (30×, 50× ), and two ONT Duplex (30×, 50× ) datasets. The read-length distributions of these sequencing datasets are consistent with real long-read sequencing data (Supplementary Table S1). We consider a shorter read as contained in a longer read if the shorter read is a proper substring of the longer read. A read is *non-contained* if it is not contained in any other read.

We tested RAFT-Hifiasm and Hifiasm methods to evaluate their ability to address the issue of assembly gaps that occur due to contained read deletion. The standard string graph formulation [21] uses non-contained reads and ignores contained reads. Hifiasm [5] uses non-contained reads to build its initial string graph and rescues a small number of contained reads later. In the RAFT-Hifiasm method, RAFT outputs a set of fragmented reads. The string graph is constructed using non-contained reads in the fragmented sequencing data. Again, Hifiasm attempts to rescue some contained reads. The benefit of using simulated data in this experiment is that we know the sampling interval of the reads in the original genome sequence. One way to spot an assembly gap due to contained read deletion is by aligning the set of reads retained in a string graph to the HG002 genome. Any interval in the genome which has zero read-alignment coverage but non-zero sequencing depth (w.r.t. the original set of reads) corresponds to an assembly gap caused by contained read deletion (Methods).

RAFT-Hifiasm outperformed Hifiasm in this experiment. Using RAFT-Hifiasm, we were able to reduce the number of assembly gaps after contained read deletion by at least an order of magnitude. We eliminated the gaps entirely in two datasets (Table 1). A small number of assembly gaps due to contained read deletion remain when using RAFT-Hifiasm because RAFT preserves repetitive regions in reads. RAFT increases the fraction of non-contained bases by narrowing the read-length read distribution. Accordingly, the fractions of bases used in RAFT-Hifiasm’s graphs are higher. We will demonstrate the impact of this approach on improving assembly quality in the next section.

**Table 1:**
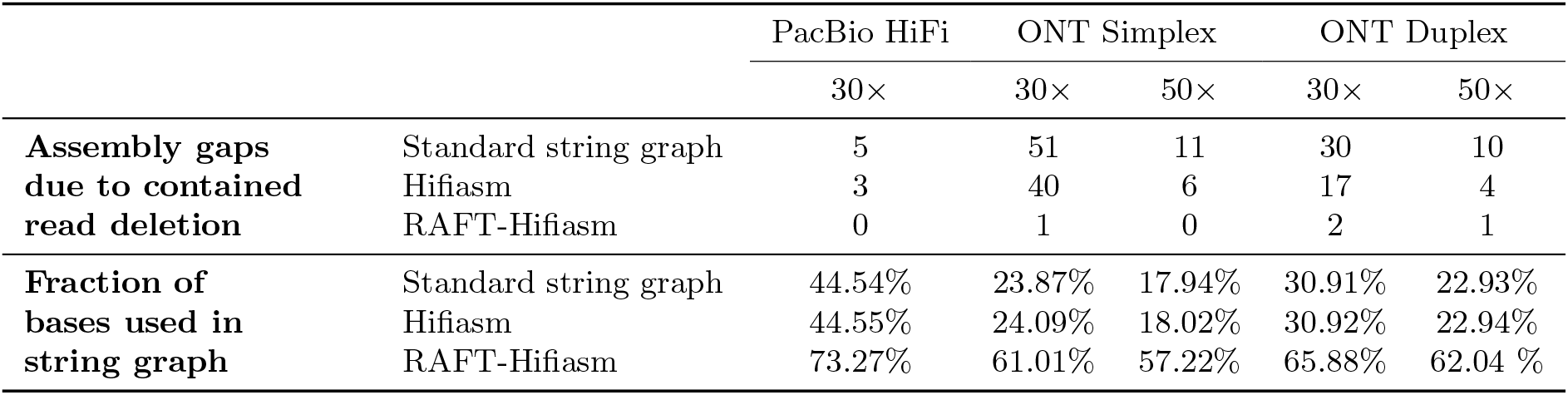
Evaluation of RAFT-HiFiasm using simulated data. We show the count of assembly gaps in the string graphs constructed by three methods, and the fraction of bases used in the graphs.

|  |  | PacBio HiFi | ONT Simplex |  | ONT Duplex |  |
| --- | --- | --- | --- | --- | --- | --- |
|  |  | 30× | 30× | 50× | 30× | 50× |
| **Assembly gaps due to contained read deletion** | Standard string graph | 5 | 51 | 11 | 30 | 10 |
|  | Hifiasm | 3 | 40 | 6 | 17 | 4 |
|  | RAFT-Hifiasm | 0 | 1 | 0 | 2 | 1 |
| **Fraction of bases used in string graph** | Standard string graph | 44.54% | 23.87% | 17.94% | 30.91% | 22.93% |
|  | Hifiasm | 44.55% | 24.09% | 18.02% | 30.92% | 22.94% |
|  | RAFT-Hifiasm | 73.27% | 61.01% | 57.22% | 65.88% | 62.04 % |

### 2.5 Evaluation using real data

We tested the RAFT-Hifiasm workflow using four publicly available real datasets comprising long and accurate reads sampled from the HG002 human genome. The first dataset, D1, comprises PacBio HiFi reads with 36 × coverage. Dataset D2 is an ONT Duplex sequencing dataset with 32× coverage. Dataset D3 is a combination of D1 and D2 datasets; thus, its coverage is 68× . Dataset D4 is a high-accuracy ultra-long ONT dataset with 40× coverage. The read-length statistics of these datasets are available in Supplementary Table S1. We assembled these datasets using RAFT-Hifiasm and Hifiasm methods to compare their output. The commands and software versions for reproducing the analysis are listed in Supplementary Note S3. We skipped comparison with other recent long-read assemblers such as Verkko [25], and LJA [3]. Both Verkko and LJA are de Bruijn graph-based assemblers and as such do not share the limitations of contained read deletion caused in the string graph-based assemblers like Hifiasm. Also, without complementary parental data or Hi-C data, neither LJA nor Verkko produces phased or partially-phased assemblies.

We expected improvements in the datasets comprising ONT reads (D2-D4) because these datasets have wide read-length distributions. The results obtained using RAFT-Hifiasm and Hifiasm methods are shown in Table 2. Applying the RAFT algorithm on datasets D2-D4 improved the assembly contiguity. The NG50 statistic is defined such that 50% of the estimated size of assembly (3.1 Gbp) is realized by contigs of NG50 length or longer. The RAFT-Hifiasm method generated more contiguous assemblies using datasets D2-D4 as indicated by the contig NG50 metric and the count of T2T-complete contigs. The switch error rate was not significantly impacted by RAFT’s read fragmentation, which suggests that Hifiasm does not face any additional difficulty phasing the fragmented reads during assembly. Compared to the assemblies produced by Hifiasm for datasets D2-D4, the assemblies obtained by RAFT-Hifiasm improved gene completeness and reduced the percentage of false duplications. Further evaluation of the assemblies, including an assessment using the Genome in a Bottle benchmark [34], is provided in Supplementary Table S2.

**Table 2:** Evaluation of the RAFT-Hifiasm workflow for computing phased genome assembly. We measured assembly quality statistics separately for both haplotypes. The contig NG50 was computed by assuming a genome length of 3.1 Gbp. The tools and commands used to measure the assembly statistics are available in Supplementary Note S3.

| Dataset | Method | Size | NG50 | Switch | T2T | Multicopy | Gene completeness |  |
| --- | --- | --- | --- | --- | --- | --- | --- | --- |
|  |  | (Gbp) | (Mbp) | error (%) | contigs | genes retained(%) | Complete (%) | Duplicated (%) |
| D1: HiFi (36×) | Hifiasm | 3.038/3.040 | 59.0/45.4 | 1.09/0.96 | 0 | 83.37/76.42 | 97.59/97.70 | 0.45/0.31 |
|  | RAFT-Hifiasm | 3.039/2.982 | 44.9/62.4 | 1.06/1.01 | 1 | 79.22/82.09 | 97.87/97.45 | 0.34/0.39 |
| D2: ONT Duplex (32×) | Hifiasm | 2.995/3.045 | 42.2/51.0 | 2.37/1.66 | 2 | 80.98/81.45 | 96.94/96.63 | 1.06/1.37 |
|  | RAFT-Hifiasm | 2.965/3.038 | 80.3/52.6 | 2.02/1.90 | 6 | 83.53/80.98 | 97.72/97.59 | 0.50/0.50 |
| D3: HiFi (36×) + ONT Duplex (32×) | Hifiasm | 3.130/2.986 | 49.3/53.3 | 0.82/1.00 | 1 | 83.85/77.78 | 95.69/95.79 | 2.42/2.10 |
|  | RAFT-Hifiasm | 3.033/3.020 | 89.6/89.3 | 0.94/1.10 | 7 | 83.13/80.50 | 97.79/97.98 | 0.42/0.41 |
| D4: ONT high-acc UL (40×) | Hifiasm | 3.439/3.435 | 16.2/20.8 | 2.49/1.69 | 0 | 74.02/78.18 | 67.72/70.41 | 19.88/19.82 |
|  | RAFT-Hifiasm | 3.047/3.072 | 81.3/49.1 | 2.19/1.87 | 1 | 81.45/77.45 | 97.21/96.94 | 0.71/0.51 |

Hifiasm used 8.5 hours to assemble the largest dataset, D3, on a CPU-based server with 128 cores. In contrast, the RAFT-Hifiasm workflow took 17.7 hours. Currently, the RAFT-Hifiasm workflow executes RAFT once and Hifiasm thrice. The three Hifiasm runs are used for long-read error-correction, computing all-to-all read alignments, and computing genome assembly, respectively. RAFT’s runtime share was only about 1.2 hours. A tighter software integration of RAFT and Hifiasm in the future may help to avoid redundant steps and optimize the runtime.

## 3. Discussion

This paper analyses and addresses a longstanding weakness of the string graph formulation [21, 22]. String graphs have been commonly used in several *de novo* genome and metagenome assemblers during the past three decades, but the issue of assembly gaps caused by contained read deletion came to limelight only recently [4, 13, 19, 23]. The quality of modern haplotype-resolved genome assemblies has improved to an extent where the few assembly gaps caused by contained read deletion are now noticeable [19]. Contained read deletion occasionally leads to the loss of useful reads in regions of low heterozygosity of diploid or polyploid genomes and in the highly repetitive regions of genomes. In both cases, all reads that support an interval in the genome may be discarded when all of them are contained in a longer read sampled from a near-identical but different region of the genome.

We presented the CGProb method to count the frequency of observing assembly gaps due to contained read deletion in a string graph. We measured the frequency for different read-length distributions and sequencing depths. This is the first mathematical model developed to assess this problem. Our analysis showed that assembly gaps due to contained read deletion are at least one order of magnitude more frequent in ONT sequencing outputs than PacBio HiFi sequencing outputs because the latter have much less variability in read lengths. In both cases, the frequency dropped rapidly with an increase in the sequencing coverage. Our method can help users to compare the relative frequencies of an assembly gap for different sequencing technologies at the same sequencing depth, or the same sequencing technology at different sequencing depths. CGProb currently works under the assumptions of error-free reads and a single heterozygous locus in the diploid genome. In future versions of CGProb, we hope to further extend the theory and relax these assumptions, e.g., to compute the frequency of assembly gaps for genome sequences that are heterozygous at more than one closely-spaced loci, or for genome sequences containing repetitive sequences, or when ploidy exceeds two. Further analysis may help to characterize the regions of a genome where assembly gaps are more likely and motivate novel methods to address the issue.

We also presented RAFT as a solution to address the issue of contained reads by fragmenting reads and obtaining a more uniform read-length distribution. Using ONT Duplex reads and a mixture of ONT Duplex and PacBio HiFi reads, combining RAFT and Hifiasm improved the assembly contiguity as evidenced by the increased contig NG50 and the number of contigs assembled T2T (Table 2). We observed significant improvements in assembly contiguity when using high-accuracy ultra-long ONT reads as well. We expect that further advances in the accuracy of ONT sequencing and haplotype-aware error-correction algorithms [26] would also make ONT Simplex reads amenable to the RAFT-Hifiasm approach. The use of ONT Simplex reads will be useful to achieve a scalable and low-cost method for generating T2T haplotype-resolved genome assemblies. Although we specifically chose to use Hifiasm alongside RAFT, the RAFT approach is easy to implement and can be slotted into any overlap-based assembly algorithm. It should work well with any read sequencing data which has a wide read length distribution and high per-base accuracy. RAFT complements another promising approach that uses ultra-long reads to rescue useful contained reads [4].

## 4 Methods

### 4.1 Counting sequencing outputs containing an assembly gap

In the following, we formally present the details of the CGProb method that calculates the fraction of sequencing outputs which contain an assembly gap in the string graph. We will first state our simplifying assumptions, define the set of valid sequencing outputs, and characterise those sequencing outputs that are affected by the deletion of contained reads. Subsequently, we will count the set of valid sequencing outputs and the affected sequencing outputs combinatorially using generating functions [32] and the inclusion-exclusion principle [1].

#### Assumptions and notations

We make a few simplifying assumptions to make the theoretical analysis tractable. We consider a genome with a single chromosome and ploidy= 2. We represent each haplotype sequence as a circular string to avoid complications arising from boundaries. Suppose both haplotype sequences have lengths *G* and differ at a single heterozygous SNP locus. Without loss of generality, we say that the heterozygous locus is at position 1 in the circular diploid genome. We assume that at least one read is sampled from the heterozygous locus on each haplotype. Each sequencing read is a substring of a haplotype sequence. Accordingly, a read is characterised by its haplotype of origin, length, and stop position. An output of a genome sequencing experiment can be represented as a pair of multisets (*S*_1_, *S*_2_), where *S*_*k*_ is a multiset of reads sampled from haplotype *k, k* = 1, 2.

We assume that repeats do not exist in our experimental setup. In other words, if a read’s sampling interval overlaps with the heterozygous locus in the genome, then the read has a unique match in its haplotype of origin and no match in the opposite haplotype. Similarly, if a read’s sampling interval does not overlap with the heterozygous locus, the read has a unique match in each haplotype. A read’s matching interval in a haplotype and all the sub-intervals of this interval are said to be *supported* by that read.

We say that read *r*_*j*_ is *contained* in read *r*_*i*_ if *r*_*j*_ is a proper substring of *r*_*i*_. For example, reads *r*_8_ and *r*_9_ are contained in read *r*_7_ in Figure 3. We use *N*_*k*_ to denote the total number of reads on haplotype *k*. Thus, the total count of reads, denoted by *N*, is *N*_1_ + *N*_2_. Let *N*_*k,i*_ denote the number of reads on haplotype *k* of length *i*. Note that Σ_*i*_ *N*_*k,i*_ = *N*_*k*_. Let, *λ*_*k*_ be the length of the longest read on haplotype *k*. We assume that, *λ*_*k*_ *< G/*2,*k* = 1, 2, i.e., the longest read length on both haplotypes is less than half the genome length. We will know the values of *N*_*k*_’s, *N*_*k,i*_’s, and, *λ*_*k*_’s from the user-specified read-length distribution and per-haplotype sequencing depths.

**Figure 3:**
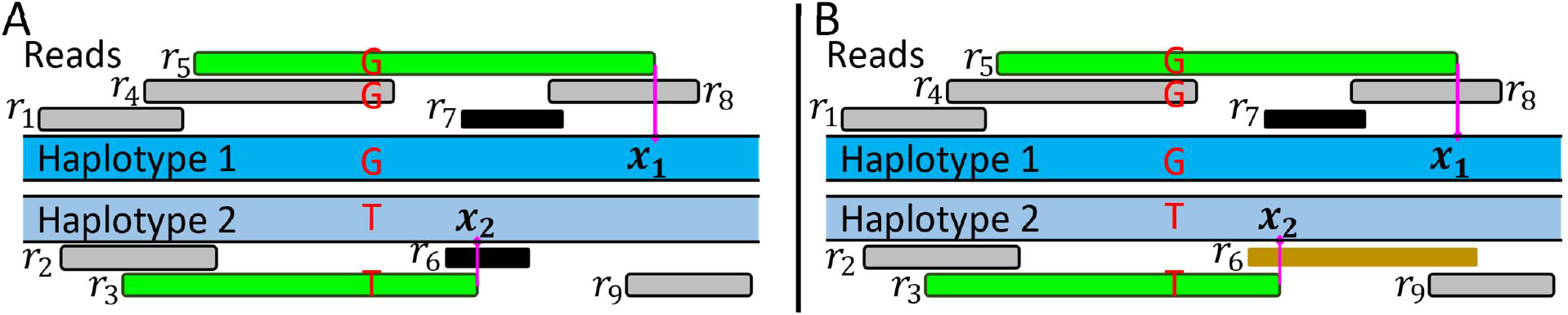
**(A)** An example of a sequencing output that is affected by the deletion of contained reads *r*_6_ and *r*_7_. Removing contained reads *r*_6_ and *r*_7_ introduces an assembly gap on haplotype 2. **(B)** An example of a sequencing output where contained read deletion does not introduce an assembly gap. Read *r*_6_ supports the sampling interval of contained read *r*_7_ after its deletion.

#### Size of the set of valid sequencing outputs

Once the values of *N*_*k,i*_’s are known, we define *S* to be the set of all valid sequencing outputs consistent with our stated assumptions. We note again that we only consider those sequencing outputs which contain at least one read sampled from each haplotype supporting the heterozygous locus, i.e., the interval [1, 1]. For that reason, we compute four quantities *T, T*_1_, *T*_2_, *T*_12_:

1. *T* is the cardinality of the set of those sequencing outputs having *N*_*k,i*_ reads of length *i* on haplotype *k* for all *i 2* [1,, *λ*_*k*_] and for all *k 2 {*1, 2*}*.
2. *T*_1_ is defined similarly to *T* but with a constraint that no read supports [1, 1] on haplotype 1.
3. *T*_2_ is defined similarly to *T* but with a constraint that no read supports [1, 1] on haplotype 2.
4. *T*_12_ is defined similarly to *T* but with a constraint that no read supports [1, 1] on either haplotype.

Using the principle of inclusion and exclusion, we have |*S*| = *T - T*_1_ *-T*_2_ + *T*_12_. We compute *T, T*_1_, *T*_2_, and *T*_12_ by writing out ordinary generating functions. We use ordinary generating functions *f*_*i,j,k*_(*x*) for reads of length *i* which stop at position *j* on haplotype *k*. The monomial *x*^*n*^ in *f*_*i,j,k*_(*x*) stands for *n* identical reads of length *i* which stop at position *j* on haplotype *k*. The coefficient of *x*^*n*^ in *f*_*i,j,k*_(*x*) is either 1 or 0, which indicates whether or not *n* identical reads having length *i* and stopping position *j* are permitted to exist. For example, the coefficient of 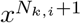 in *f*_*i,j,k*_(*x*) is 0 because *N*_*k,i*_ + 1 reads of length *i* don’t exist on haplotype *k*. The number of multisets of reads of length *i* on haplotype *k*, denoted by *ff*_*k,i*_, is the coefficient of 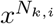 in the product 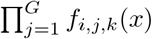. To obtain the total number of multisets of sequencing outputs, we compute 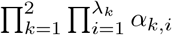.

To estimate *T*, we set the ordinary generating functions of reads of length *i* stopping at position *j* on haplotype *k* to 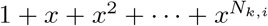, for *j* = 1 to *G*. This is because we don’t restrict the existence of any reads in this case. When estimating *T*_1_, we set the ordinary generating functions of reads of length *i* stopping at position *j* on haplotype *k* to be 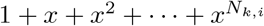 provided the read doesn’t support the interval [1, 1] on haplotype 1. If it does, then that ordinary generating function is the polynomial *x*^0^, because no such reads are permitted. Similarly, for estimating *T*_2_, we restrict reads on haplotype 2 from supporting [1, 1]. Lastly, for *T*_12_, we set the ordinary generating functions of all reads supporting [1, 1] to *x*^0^.

#### Assembly gap due to contained read deletion

Next, we formally define the occurrence of an assembly gap in a string graph due to contained read deletion. Among all reads on haplotype 1 which support the interval [1, 1], let *x*_1_ be the stop position of those reads which stop closest to position,*\*_1_. Similarly, let *x*_2_ be the stop position of the reads which support [1, 1] and stop closest to position, *λ* on haplotype 2. These reads are shown in green in Figure 3. *x*_1_ and *x*_2_ are well-defined for a given sequencing output. Having identified *x*_1_ and *x*_2_ for a sequencing output ℛ, we say that ℛ belongs to class (*x*_1_, *x*_2_). Therefore, this assignment partitions the set of all valid sequencing outputs.

##### Definition 1.

*A sequencing output belonging to class* (*x*_1_, *x*_2_) *is said to have an* ***assembly gap due to contained read deletion*** *if the interval* [min(*x*_1_, *x*_2_), min(*x*_1_, *x*_2_) + 1] *is originally supported on both haplotypes by some reads, and no longer supported on at least one haplotype by any read after the deletion of contained reads*.

Figure 3A shows an example of a sequencing output where *x*_1_ *> x*_2_ and the deletion of contained reads leads to the loss of reads supporting the interval [*x*_2_, *x*_2_ + 1] on haplotype 2. Other gaps in the assembly may also occur naturally due to a lack of coverage on a region in the original sequencing data. These gaps are not considered in our analysis because these are not introduced computationally. On the other hand, an assembly gap in Definition 1 is artificially introduced by a graph simplification heuristic and hinders an assembler from phasing the heterozygous variant. Using Definition 1, we restrict our attention to the interval [min(*x*_1_, *x*_2_), min(*x*_1_, *x*_2_) + 1], which is located on the clockwise side of the heterozygous locus in the circular genome. Next, we establish the distinguishing property of the sequencing outputs that are affected by contained read deletion. We will use this property for counting these sequencing outputs.

##### Lemma 1.

*Let λ be a sequencing output belonging to class* (*x*_1_, *x*_2_). *λ has an assembly gap due to contained read deletion if and only if λ satisfies all the following conditions: (1) x*_1_ *6*= *x*_2_, *(2) At least one read starting in* [2, min(*x*_1_, *x*_2_)] *on either haplotype stops in* [1 + min(*x*_1_, *x*_2_), max(*x*_1_, *x*_2_)], *and (3) No read starting in* [2, min(*x*_1_, *x*_2_)] *on either haplotype supports* [1 + max(*x*_1_, *x*_2_), 1 + max(*x*_1_, *x*_2_)].

*Proof*. Let *ℛ* be a sequencing output which satisfies the three conditions. Without loss of generality, assume *x*_1_ *> x*_2_. Let *𝒳* be the multiset of reads that start in [2, *x*_2_] and stop in [1 + *x*_2_, *x*_1_] on either haplotype. Each read in *𝒳* supports the interval [*x*_2_, *x*_2_ + 1] on both haplotypes. The second condition guarantees that |*𝒳* | *2’* 1. However, all reads in *𝒳* are contained in some read that supports the interval [1, *x*_1_] on haplotype 1. Accordingly, all reads in *𝒳* will be removed by the contained read deletion heuristic. The third condition ensures that no other read in ℛ supports the interval [*x*_2_, *x*_2_ + 1] on haplotype 2. As a result, an assembly gap due to contained read deletion is guaranteed.

Conversely, suppose *ℛ* is a sequencing output which fails to satisfy one of the three conditions. In each case, we prove that an assembly gap due to contained read deletion does not occur in the string graph of *ℛ*.

##### Condition (1)

Suppose *ℛ* fails to satisfy the first condition. Therefore, *x*_1_ = *x*_2_. In this case, min(*x*_1_, *x*_2_) = max(*x*_1_, *x*_2_) = *x*_1_. If no read in *ℛ* supports [*x*_1_, *x*_1_ + 1] on haplotype 1, then by Definition 1, *ℛ* cannot have an assembly gap due to contained read deletion. Accordingly, let us consider the non-empty multiset *𝒴* of the reads that support [*x*_1_, *x*_1_ + 1] on haplotype 1. Let *r* be a read with the maximum length in *𝒴*. By the definition of *x*_1_ and *x*_2_, read *r* cannot be contained in a read which supports [1, 1] on haplotype 2. For that reason, any read containing *r* must support [*x*_1_, *x*_1_ + 1] on haplotype 1. Such a read cannot exist because we selected *r* with the maximum length from *𝒴*. Thus, after deleting all the contained reads, read *r* will support [*x*_1_, *x*_1_ + 1] on both haplotypes.

##### Condition (2)

Suppose *ℛ* satisfies the first condition and does not satisfy the second. Without loss of generality, assume *x*_1_ *> x*_2_. We know that no reads start in [2, *x*_2_] on either haplotype and stop in [1+*x*_2_, *x*_1_]. Let us analyse the reads supporting the interval [*x*_2_, *x*_2_ + 1]. Case (a): There is no read in *ℛ* which supports [*x*_2_, *x*_2_ + 1] on haplotype 2 before the deletion of contained reads. Then, *ℛ* does not have an assembly gap due to contained read deletion. Case (b): One or more reads in *ℛ* support [*x*_2_, *x*_2_ + 1] on haplotype 2. Then we must have a read in ℛ starting in [2, *x*_2_] and supporting [1 + *x*_1_, 1 + *x*_1_] on both haplotypes. Let *r* be a read with the maximum length satisfying this condition. Read *r* cannot be a contained read for the same reason as stated earlier. Thus, *r* continues to support [*x*_2_, *x*_2_ + 1] on both haplotypes after the deletion of contained reads does not introduce an assembly gap in *ℛ*.

##### Condition (3)

Suppose *ℛ* satisfies the first and second conditions, and fails to satisfy the third. Without loss of generality, assume *x*_1_ *> x*_2_. Failure to satisfy condition (3) means that there exist one or more reads *∈ ℛ* which start in [2, *x*_2_] and support [*x*_1_ + 1, *x*_1_ + 1] on both haplotypes. Arguing along the lines of Case (b) of Condition (2), consider a longest read which starts in [2, *x*_2_] and supports [*x*_1_ + 1, *x*_1_ + 1]. This is not a contained read and will continue to support [*x*_2_, *x*_2_ + 1] after the deletion of contained reads. This implies that an assembly gap due to contained read deletion does not occur in *ℛ*.

We will denote *M* as the set of sequencing outputs containing an assembly gap due to contained read deletion. The ratio 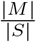 will give us the fraction of sequencing outputs with a user-specified read length distribution containing an assembly gap due to contained read deletion. Let us denote the set of sequencing outputs that belong to class (*x*_1_, *x*_2_) and satisfy Lemma 1 as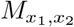 . We describe our method for calculating 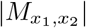 in Supplementary Note S1. Using this method, we calculate 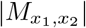 for all *x*_1_ *2* [1,, *λ*_1_] and *x*_2_ *∈* [1,, *λ*] and add these to obtain |*M*|.

##### Lemma 2.

*The total number of sequencing outputs containing assembly gaps due to contained read deletion is* 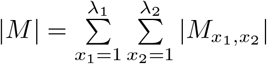.

We implemented the methods for calculating |*M*| and |*S*| in CGProb. The proposed approach is significantly more efficient when compared to a naive method of individually analysing *O*(*G*^*N*^ ) sequencing outputs. The time complexity of our method is polynomially bounded.

#### CGProb implementation details and experimental setup

We set genome length *G* to 1 Mbp. CGProb condenses the genome length and the lengths of each read by a user-specified factor (default value = 1000) to speed up the computation. We used arbitrary-precision integer arithmetic [10] to eliminate numerical error. The read-length distribution can be extracted as a list of distinct read lengths and counts from any long-read sequencing experiment. In our experiments (Section 2.2), we used three read-length distributions corresponding to PacBio HiFi, ONT Simplex, and ONT Duplex technologies. We obtained these distributions using publicly available datasets. Using the list of distinct read lengths and counts, the value of *G*, and the per-haplotype sequencing depths, we ran Seqrequester (commit: `31141c1`) [30] five times for each input. Each run was used to simulate a revised list of read lengths and counts on each haplotype. CGProb’s runtime was 2 hours on a 50x HiFi dataset, 2 hours on a 50x ONT Duplex dataset, and 15 hours on a 50x ONT Simplex dataset on a server with two 24-core Intel Xeon Gold 6248R CPUs.

### 4.2 RAFT implementation and benchmarking

#### RAFT implementation details

In the RAFT-Hifiasm workflow, RAFT uses error-corrected reads, and all-to-all read alignments computed by Hifiasm. In our experiments, we ran Hifiasm (v0.19.8-r603) two times; the first run executed with `--write-ec` parameter returned the error-corrected reads, and the second run executed with `--dbg-ovec` parameter returned all-to-all alignments in the pairwise alignment format (PAF). RAFT places *potential breakpoint markers* on each read. These markers are positioned evenly at intervals of 5 kbp by default. In the highly-repetitive regions of the genome, it is useful to retain long reads to avoid contig breaks. Accordingly, RAFT deletes the markers at those bases which are predicted to be sampled from repetitive regions of the genome. RAFT uses all-to-all alignments to predict repetitive bases in a read. While processing read *r*, it counts the number of overlapping reads on each base of read *r*. Assuming *cov* is the sequencing depth of the input sequencing dataset, and *μ* is a cutoff parameter (default value = 1.5), RAFT identifies all intervals of length *2’* 5 kbp in read *r*, where the count exceeds *μ · cov* for all bases of the interval. RAFT further extends these intervals by 500 bp on both sides to avoid having a marker very close to a repeat. After deleting the markers from these intervals, RAFT uses the remaining markers for fragmenting the read into shorter reads. The length of fragmented reads in RAFT is set to 20 kbp as default. Starting from the first base of the read, it finds the first marker located after the first 20k non-repetitive bases and breaks the read at the marker. Similarly, it finds the first marker after the subsequent 20k non-repetitive bases of the read and cuts again. The process continues until it reaches the last base of the read. The user can adjust the algorithm parameters using RAFT’s command line interface.

#### Simulation-based benchmarking procedure

We evaluated the count of coverage gaps caused by contained read deletion using the standard string graph formulation [21], Hifiasm (v0.19.8-r603) [5], and RAFT-Hifiasm. As discussed earlier, Hifiasm rescues a small number of contained reads after building a string graph. The RAFT-Hifiasm method follows a different approach by fragmenting the input reads. We benchmarked the three methods by sampling error-free long-reads from Hifiasm’s trio-based assembly of HG002 human genome. We generated five sets of reads using Seqrequester, resembling the read-length distributions of real PacBio HiFi, ONT Simplex, and ONT Duplex sequencing datasets. Given a read set, we identified the set of non-contained reads by generating all-to-all read overlaps using Minimap2 (v2.26-r1175) [18]. First, we identified contained reads entirely encompassed within a longer read with 100% alignment identity based on the Minimap2 output. Accordingly, the set of non-contained reads comprised all reads which were not contained.

To evaluate the standard string graph formulation, we aligned the set of non-contained reads to the two HG002 haplotype assemblies, one at a time. We used Bedtools (v2.29.1) [24] to extract all genomic intervals with zero alignment coverage. Next, we excluded the intervals that overlapped with any interval with zero sequencing depth. We also excluded the intervals which included the first or last 25 kbp bases of a HG002 contig to account for edge effects. This left us with a subset of intervals that accurately represented the assembly gaps caused by contained read deletion. We followed the same procedure for Hifiasm but modified the last step because Hifiasm rescues a few contained reads. We ran Hifiasm to compute the rescued reads and aligned the rescued reads to the two HG002 haplotype assemblies. We reported the assembly gaps which remained unresolved. Our benchmarking procedure for RAFT-Hifiasm was similar to Hifiasm, except we considered the set of fragmented reads produced by RAFT instead of the original simulated reads. We recomputed the sets of non-contained reads and rescued reads by following the same procedure.

#### Computing assembly quality statistics

We followed Hifiasm’s benchmarking procedure [5, 6] to assess assembly quality. We evaluated the phased assemblies (Table 2, Supplementary Table S2) computed by Hifiasm (v0.19.8-r603) and RAFT-Hifiasm methods. A phased assembly comprises two sets of contigs from haplotypes 1 and 2, respectively. In Table 2, we estimated NG50 using QUAST (v5.2.0) [11] by assuming the size of true assembly as 3.1 Gbp. Additionally, we used: (i) Asmgene [18] to compute gene completeness, (ii) Yak (v0.1-r69-dirty) with parental sequencing data to estimate the switch error rate, and (iii) HPRC workflow (https://github.com/biomonika/HPP/blob/main/assembly/wdl/workflows/assessAsemblyCompletness.wdl) to estimate the count of T2T-complete contigs. The commands used to run these tools are available in Supplementary Note S3.

## Data availability

We used publicly available datasets to obtain read length distributions for the analysis using CGProb. For the PacBio HiFi read length distribution, we used data from SRR10382244, SRR10382245, SRR10382248, and SRR10382249 from the SRA. We obtained the ONT Simplex read length distribution from (https://s3-us-west-2.amazonaws.com/human-pangenomics/index.html?prefix=working/HPRC_PLUS/HG02080/raw_data/nanopore/HG02080_2.fastq.gz), and ONT Duplex read length distribution from (https://human-pangenomics.s3.amazonaws.com/index.html?prefix=submissions/0CB931D5-AE0C-4187-8BD8-B3A9C9BFDADE--UCSC_HG002_R1041_Duplex_Dorado/Dorado_v0.1.1/stereo_duplex/*_stereo_duplex_pass.fastq.gz) datasets.

For the simulation-based benchmarking procedure, we used a Trio-based assembly of the HG002 human genome by Hifiasm (ftp://ftp.dfci.harvard.edu/pub/hli/hifiasm-phase/v2/HG002.hifiasm.trio.0.16.1.hap1.fa.gz, ftp://ftp.dfci.harvard.edu/pub/hli/hifiasm-phase/v2/HG002.hifiasm.trio.0.16.1.hap2.fa.gz).

For benchmarking the RAFT-Hifiasm workflow, we used four datasets. Dataset D1 contained PacBio Hifi reads from SRR10382244, SRR10382245, SRR10382248, and SRR10382249. Dataset D2 contained ONT Duplex reads from https://human-pangenomics.s3.amazonaws.com/index.html?prefix=submissions/0CB931D5-AE0C-4187-8BD8-B3A9C9BFDADE--UCSC_HG002_R1041_Duplex_Dorado/Dorado_v0.1.1/stereo_duplex/*_stereo_duplex_pass.fastq.gz. Dataset D4 contained ONT high-accuracy ultra-long reads from https://labs.epi2me.io/gm24385_ncm23_preview/.

To measure switch error rate in the assemblies, we used complementary Illumina parental data from HG003 and HG004 samples located at (https://s3-us-west-2.amazonaws.com/human-pangenomics/index.html?prefix=NHGRI_UCSC_panel/HG002/hpp_HG002_NA24385_son_v1/parents/ILMN/downsampled/HG003/ and https://s3-us-west-2.amazonaws.com/human-pangenomics/index.html?prefix=NHGRI_UCSC_panel/HG002/hpp_HG002_NA24385_son_v1/parents/ILMN/downsampled/HG004/. Gene completeness metrics were computed using asmgene with cDNA data obtained from http://ftp.ensembl.org/pub/release-102/fasta/homo_sapiens/cdna/Homo_sapiens.GRCh38.cdna.all.fa.gz. For computing assembly QV using Yak (Supplementary Table S2), we used Hi-C data files ‘HG002.HiC_1*.fastq.gz’ from https://github.com/human-pangenomics/HG002_Data_Freeze_v1.0. GIAB small variant benchmark v4.2.1 was obtained from https://ftp-trace.ncbi.nlm.nih.gov/ReferenceSamples/giab/release/AshkenazimTrio/HG002_NA24385_son/NISTv4.2.1/GRCh38/ for evaluating the variant calls obtained using genome assemblies.

## Code availability

CGProb is available as an open-source tool on GitHub (https://github.com/at-cg/CGProb), commit ID: `0f16cfc`. RAFT is available as an open-source tool on GitHub (https://github.com/at-cg/RAFT), commit ID: `f1e34ad`. Other tools used in this study and their software versions are provided in Supplementary Table S3.

## Competing interest statement

The authors declare no competing interests.

## Acknowledgements

The authors thank Mile Sikic, Sunil Chandran for providing useful feedback. Josipa Lipovac and Prasad Sarashetti tested RAFT code and shared valuable feedback. Haoyu Cheng addressed our concerns regarding Hifiasm. We thank the Human Pangenome Reference Consortium for making their sequencing datasets publicly available. This research is supported in part by the funding from the DBT/Wellcome Trust India Alliance (IA/I/23/2/506979) and the National Supercomputing Mission, India under DST/NSM/R&D HPC Applications. We also thank the Council of Scientific and Industrial Research (CSIR), Ministry of Science and Technology, India, for the financial support through the Junior Research Fellowship. We used computing resources provided by the National Energy Research Scientific Computing Center (NERSC), USA.

## Footnotes

1 Variant allele frequency is the fraction of reads supporting a specific DNA variant divided by the overall coverage at that locus.

## Notes

### Competing Interest Statement

The authors have declared no competing interest.

### Summary of Updates

Section on Results updated to reflect experimental results from updated software. CGProb performs all computations with arbitrary precision integers. Figure 1 updated. Table 2 includes results from a high-accuracy ultralong ONT dataset. Section on Methods updated with characterising criterion for assembly gaps due to contained read deletion. Data and Code availability sections added to display locations of data and code used in one location.

## References

[1] Reginald BJT Allenby and Alan Slomson. How to count: An introduction to combinatorics. CRC Press, 2010.

[2] Jasmijn A Baaijens, Amal Zine El Aabidine, Eric Rivals, and Alexander Schönhuth. “De novo assembly of viral quasispecies using overlap graphs”. In: Genome research 27.5 (2017), pp. 835–848.

[3] Anton Bankevich, Andrey V Bzikadze, Mikhail Kolmogorov, Dmitry Antipov, and Pavel A Pevzner. “Multiplex de Bruijn graphs enable genome assembly from long, high-fidelity reads”. In: Nature biotechnology 40.7 (2022), pp. 1075–1081.

[4] Haoyu Cheng, Mobin Asri, Julian Lucas, Sergey Koren, and Heng Li. “Scalable telomere-to-telomere assembly for diploid and polyploid genomes with double graph”. In: arXiv preprint 2306.03399 (2023).

[5] Haoyu Cheng, Gregory T Concepcion, Xiaowen Feng, Haowen Zhang, and Heng Li. “Haplotype-resolved de novo assembly using phased assembly graphs with hifiasm”. In: Nature methods 18.2 (2021), pp. 170–175.

[6] Haoyu Cheng, Erich D Jarvis, Olivier Fedrigo, Klaus-Peter Koepfli, Lara Urban, Neil J Gemmell, and Heng Li. “Haplotype-resolved assembly of diploid genomes without parental data”. In: Nature Biotechnology 40.9 (2022), pp. 1332–1335.

[7] Chen-Shan Chin, David H Alexander, Patrick Marks, Aaron A Klammer, James Drake, Cheryl Heiner, Alicia Clum, Alex Copeland, John Huddleston, Evan E Eichler, et al. “Nonhybrid, finished microbial genome assemblies from long-read SMRT sequencing data”. In: Nature methods 10.6 (2013), pp. 563–569.

[8] Chen-Shan Chin, Paul Peluso, Fritz J Sedlazeck, Maria Nattestad, Gregory T Concepcion, Alicia Clum, Christopher Dunn, Ronan O’Malley, Rosa Figueroa-Balderas, Abraham Morales-Cruz, et al. “Phased diploid genome assembly with single-molecule real-time sequencing”. In: Nature methods 13.12 (2016), pp. 1050–1054.

[9] Xiaowen Feng, Haoyu Cheng, Daniel Portik, and Heng Li. “Metagenome assembly of high-fidelity long reads with hifiasm-meta”. In: Nature Methods 19.6 (2022), pp. 671–674.

[10] Torbjörn Granlund. “The GNU multiple precision arithmetic library”. In: http://gmplib.org/ (2010).

[11] Alexey Gurevich, Vladislav Saveliev, Nikolay Vyahhi, and Glenn Tesler. “QUAST: quality assessment tool for genome assemblies”. In: Bioinformatics 29.8 (2013), pp. 1072–1075.

[12] Joseph Hui, Ilan Shomorony, Kannan Ramchandran, and Thomas A. Courtade. “Overlap-based genome assembly from variable-length reads”. In: 2016 IEEE International Symposium on Information Theory (ISIT). 2016, pp. 1018–1022. doi: 10.1109/ISIT.2016.7541453.

[13] Chirag Jain. “Coverage-preserving sparsification of overlap graphs for long-read assembly”. In: Bioinformatics 39.3 (2023), btad124.

[14] Erich D Jarvis, Giulio Formenti, Arang Rhie, Andrea Guarracino, Chentao Yang, Jonathan Wood, Alan Tracey, Francoise Thibaud-Nissen, Mitchell R Vollger, David Porubsky, et al. “Semi-automated assembly of high-quality diploid human reference genomes”. In: Nature 611.7936 (2022), pp. 519–531.

[15] Sergey Koren, Brian P Walenz, Konstantin Berlin, Jason R Miller, Nicholas H Bergman, and Adam M Phillippy. “Canu: scalable and accurate long-read assembly via adaptive k-mer weighting and repeat separation”. In: Genome research 27.5 (2017), pp. 722–736.

[16] Heng Li. Concepts in phased assemblies. https://lh3.github.io/2021/04/17/concepts-in-phased-assemblies. 2021.

[17] Heng Li. “Minimap and miniasm: fast mapping and de novo assembly for noisy long sequences”. In: Bioinformatics 32.14 (2016), pp. 2103–2110.

[18] Heng Li. “Minimap2: pairwise alignment for nucleotide sequences”. In: Bioinformatics 34.18 (2018), pp. 3094–3100.

[19] Heng Li and Richard Durbin. “Genome assembly in the telomere-to-telomere era”. In: ArXiv (2023).

[20] Glennis A Logsdon, Mitchell R Vollger, and Evan E Eichler. “Long-read human genome sequencing and its applications”. In: Nature Reviews Genetics 21.10 (2020), pp. 597–614.

[21] Eugene W Myers. “The fragment assembly string graph”. In: Bioinformatics 21.suppl 2 (2005), pp. ii79–ii85.

[22] Eugene W Myers. “Toward simplifying and accurately formulating fragment assembly”. In: Journal of Computational Biology 2.2 (1995), pp. 275–290.

[23] Sergey Nurk, Sergey Koren, Arang Rhie, Mikko Rautiainen, Andrey V Bzikadze, Alla Mikheenko, Mitchell R Vollger, Nicolas Altemose, Lev Uralsky, Ariel Gershman, et al. “The complete sequence of a human genome”. In: Science 376.6588 (2022), pp. 44–53.

[24] Aaron R Quinlan and Ira M Hall. “BEDTools: a flexible suite of utilities for comparing genomic features”. In: Bioinformatics 26.6 (2010), pp. 841–842.

[25] Mikko Rautiainen, Sergey Nurk, Brian P Walenz, Glennis A Logsdon, David Porubsky, Arang Rhie, Evan E Eichler, Adam M Phillippy, and Sergey Koren. “Telomere-to-telomere assembly of diploid chromosomes with Verkko”. In: Nature Biotechnology (2023), pp. 1–9.

[26] Dominik Stanojevic. HERRO (Haplotype-aware error correction). https://github.com/lbcb-sci/herro. 2024.

[27] Alexandru I Tomescu and Paul Medvedev. “Safe and complete contig assembly through omnitigs”. In: Journal of computational biology 24.6 (2017), pp. 590–602.

[28] Robert Vaser and Mile Šikíc. “Time- and memory-efficient genome assembly with Raven”. In: Nature Computational Science 1.5 (2021), pp. 332–336.

[29] Riccardo Vicedomini, Christopher Quince, Aaron E Darling, and Rayan Chikhi. “Strainberry: automated strain separation in low-complexity metagenomes using long reads”. In: Nature Communications 12.1 (2021), p. 4485.

[30] Brian Walenz. Seqrequester. https://github.com/marbl/seqrequester. 2023.

[31] Aaron M Wenger, Paul Peluso, William J Rowell, Pi-Chuan Chang, Richard J Hall, Gregory T Concepcion, Jana Ebler, Arkarachai Fungtammasan, Alexey Kolesnikov, Nathan D Olson, et al. “Accurate circular consensus long-read sequencing improves variant detection and assembly of a human genome”. In: Nature biotechnology 37.10 (2019), pp. 1155–1162.

[32] Herbert S Wilf. generatingfunctionology. CRC press, 2005.

[33] Chentao Yang, Yang Zhou, Yanni Song, Dongya Wu, Yan Zeng, Lei Nie, Panhong Liu, Shilong Zhang, Guangji Chen, Jinjin Xu, et al. “The complete and fully-phased diploid genome of a male Han Chinese”. In: Cell Research (2023), pp. 1–17.

[34] Justin M Zook, Nancy F Hansen, Nathan D Olson, Lesley Chapman, James C Mullikin, Chunlin Xiao, Stephen Sherry, Sergey Koren, Adam M Phillippy, Paul C Boutros, et al. “A robust benchmark for detection of germline large deletions and insertions”. In: Nature biotechnology 38.11 (2020), pp. 1347–1355.

