## Supplementary Information for "Telomere-to-telomere assembly by preserving contained reads"

---

### Content summary


- **Note S1:** Additional details of CGProb method.
- **Note S2:** Software commands for CGProb.
- **Note S3:** Software commands for RAFT.
- **Figure S1:** Frequency of observing an assembly gap near a germline heterozygous locus.
- **Figure S2:** Frequency of observing an assembly gap near a somatic heterozygous locus.
- **Figure S3:** Read length distributions corresponding to the different read sequencing technologies.
- **Table S1:** Read length statistics of sequencing datasets used for benchmarking RAFT algorithm.
- **Table S2:** Additional quality statistics of the HG002 phased assemblies produced by RAFT-Hifiasm and Hifiasm.
- **Table S3:** Software versions.


---

### Supplementary Note S1: Additional details of CGProb method

**Read groups.** For a sequencing output belonging to class $(x_1, x_2)$ and having an assembly gap due to contained read deletion, we define the following groups to partition the reads sampled from haplotype $k$ $(k = 1, 2)$. We will see later that this classification makes it convenient to enumerate all valid multisets of reads for calculating $|M_{x_1,x_2}|$.

1. Reads in group 1 span the heterozygous locus.
   - Reads in group 1(a) stop before $x_k$.
   - Reads in group 1(b) stop at $x_k$.
   - Reads in group 1(c) stop after $x_k$.
2. The sampling interval of reads in group 2 is a sub-interval of $[2, \max(x_1, x_2)]$.
   - Reads in group 2(a) start and stop in $[2, \min(x_1, x_2)]$.
   - Reads in group 2(b) start in $[2, \min(x_1, x_2)]$ and stop in $[\min(x_1, x_2) + 1, \max(x_1, x_2)]$.
   - Reads in group 2(c) start and stop in $[\min(x_1, x_2) + 1, \max(x_1, x_2)]$.
3. Reads in group 3 stop in $[\max(x_1, x_2) + 1, G]$.
   - Reads in group 3(a) start in $[2, \min(x_1, x_2)]$.
   - Reads in group 3(b) start in $[\min(x_1, x_2) + 1, G]$.

For example, the green-coloured reads in Figure 3 in the main text are in group 1(b), and the black-coloured reads are in group 2(b). Next, we show that these read groups are mutually exclusive and exhaustive. By our definitions of $x_1$ and $x_2$ in Section 4.1, group 1(c) must be empty. By Lemma 1, group 3(a) must be empty.

**Lemma 3.** *For all the sequencing outputs belonging to class $(x_1, x_2)$ which have an assembly gap due to contained read deletion, the read groups 1(a), 1(b), 2(a), 2(b), 2(c), and 3(b) are mutually exclusive and exhaustive.*

*Proof.* If the sequencing output belongs to class $(x_1, x_2)$, then by Lemma 1, $x_1 \neq x_2$. Without loss of generality, assume $x_1 > x_2$. First, we argue that the groups are mutually exclusive. We only consider reads sampled from haplotype 1. Similar arguments can be made for reads sampled from haplotype 2.

1. Reads in group 1 support the heterozygous locus. They cannot belong to groups 2 or 3. Reads in group 1(a) stop before $x_1$ and reads in group 1(b) stop at $x_1$.
2. Reads in group 2 start and stop in the interval $[2, x_1]$. Therefore, they cannot belong to group 3. Reads in group 2 have three choices – they start and stop in $[2, x_2]$, i.e. group 2(a), they start and stop in $[x_2 + 1, x_1]$, i.e. group 2(c), or they start in $[2, x_2]$ and stop in $[x_2 + 1, x_1]$, i.e. group 2(b).

Thus, all the groups are mutually exclusive, and no read can belong to two groups simultaneously.

Next, we show that reads from any sequencing output belonging to class $(x_1, x_2)$ can be classified into one of the six groups. Again, we restrict our attention to reads arising from haplotype 1.

1. All the reads which support $[1, 1]$ will be classified into groups 1(a) or 1(b), according to their stop position before $x_1$ or being $x_1$, respectively.
2. Suppose a read starts and stops in the interval $[2, x_1]$. This read will be classified into group 2 because groups 2(a), 2(b) and 2(c) include all possible sub-intervals of $[2, x_1]$.
3. The remaining reads stop in $[x_1 + 1, G]$. By Lemma 1, these reads must start in $[x_2 + 1, G]$. Hence these reads must be in group 3(b).

In the above argument, we first exhaust all reads that support $[1, 1]$. The remaining reads must start and stop in $[2, G]$. Next, we consider all stop positions in $[2, G]$. For every stop position in $[2, G]$, we have considered all valid start positions. Therefore, no read is left unclassified into a group. Thus, all reads sampled from haplotype 1 are partitioned into the six groups. □

**Counting $|M_{x_1,x_2}|$.** While counting read sequencing outputs in class $(x_1, x_2)$ with an assembly gap due to contained read deletion, we need to consider the ones which have at least one read in group 1(b) on both haplotypes, at least one read in either haplotype on group 2(b), and zero reads in group 1(c) and 3(a) on both haplotypes (Lemma 1). We use an inclusion-exclusion argument to count such outputs. To this end, we define $M_{11}, M_{12}, M_{13}, M_{14}$ as following:

- $M_{11}$ = Set of multisets of reads on haplotype 1 where groups 1(c) and 3(a) have zero reads
- $M_{12}$ = Set of multisets of reads on haplotype 1 where groups 1(b), 1(c), and 3(a) have zero reads
- $M_{13}$ = Set of multisets of reads on haplotype 1 where groups 1(c), 2(b), and 3(a) have zero reads
- $M_{14}$ = Set of multisets of reads on haplotype 1 where groups 1(b), 1(c), 2(b), and 3(a) have zero reads

Similarly, we denote the corresponding sets of multisets on haplotype 2 using symbols $M_{21}, M_{22}, M_{23}$, $M_{24}$.

**Lemma 4.** *The total number of sequencing outputs that belong to class $(x_1, x_2)$ and have an assembly gap due to contained read deletion is $|M_{x_1,x_2}| = (|M_{11}| - |M_{12}|)(|M_{21}| - |M_{22}|) - (|M_{13}| - |M_{14}|)(|M_{23}| - |M_{24}|)$.*

*Proof.* We want to count sequencing outputs which have (i) at least one read in group 1(b) from haplotype 1, (ii) at least one read in group 1(b) from haplotype 2, (iii) at least one read in group 2(b) from either haplotype 1 or 2, and (iv) exactly zero reads in group 1(c) and 3(a) on both haplotypes. We describe these sequencing outputs in terms of multisets of reads from each haplotype as follows:

(1) Multisets of reads from haplotype 1 with at least one read in group 1(b), at least one read in group 2(b), and zero reads in groups 1(c) and 3(a), taken together with multisets of reads from haplotype 2 with at least one read in group 1(b), at least one read in group 2(b), and zero reads in groups 1(c) and 3(a).
(2) Multisets of reads from haplotype 1 with at least one read in group 1(b) and exactly zero reads in groups 1(c), 2(b) and 3(a), taken together with multisets of reads from haplotype 2 with at least one read in group 1(b), at least one read in group 2(b), and zero reads in groups 1(c) and 3(a).
(3) Multisets of reads from haplotype 1 with at least one read in group 1(b), at least one read in group 2(b), and zero reads in groups 1(c) and 3(a), taken together with multisets of reads from haplotype 2 with at least one read in group 1(b) and exactly zero reads in groups 1(c), 2(b) and 3(a).

Note that these three partition the set of sequencing outputs that must be counted. Next, the following facts follow from the principle of inclusion and exclusion:

- Multisets of reads from haplotype 1 with at least one read in group 1(b), at least one read in group 2(b), and zero reads in groups 1(c) and 3(a) = $|M_{11}| - |M_{12}| - |M_{13}| + |M_{14}|$.
- Multisets of reads from haplotype 2 with at least one read in group 1(b), at least one read in group 2(b), and zero reads in groups 1(c) and 3(a) = $|M_{21}| - |M_{22}| - |M_{23}| + |M_{24}|$.
- Multisets of reads from haplotype 1 with at least one read in group 1(b) and exactly zero reads in groups 1(c), 2(b) and 3(a) = $|M_{13}| - |M_{14}|$.
- Multisets of reads from haplotype 2 with at least one read in group 1(b) and exactly zero reads in groups 1(c), 2(b) and 3(a) = $|M_{23}| - |M_{24}|$.

Thus, the number of elements in (1) is $(|M_{11}| - |M_{12}| - |M_{13}| + |M_{14}|)(|M_{21}| - |M_{22}| - |M_{23}| + |M_{24}|)$. The number of elements in (2) is $(|M_{13}| - |M_{14}|)(|M_{21}| - |M_{22}| - |M_{23}| + |M_{24}|)$. The number of elements in (3) is $(|M_{11}| - |M_{12}| - |M_{13}| + |M_{14}|)(|M_{23}| - |M_{24}|)$. Adding these gives $|M_{x_1,x_2}| = (|M_{11}| - |M_{12}|)(|M_{21}| - |M_{22}|) - (|M_{13}| - |M_{14}|)(|M_{23}| - |M_{24}|)$. □

The eight quantities $|M_{11}|, |M_{12}|, |M_{13}|, |M_{14}|, |M_{21}|, |M_{22}|, |M_{23}|, |M_{24}|$ can be efficiently calculated using ordinary generating functions [32]. First, we write an ordinary generating function $f_{i,j,k}(x)$ representing the number of valid reads of length $i$ stopping at position $j$ on haplotype $k$. Second, the coefficient of the monomial of degree $N_{k,i}$ in the product $\prod_{j=1}^{G} f_{i,j,k}(x)$ gives the number of multisets of reads of length $i$ on haplotype $k$. Third, the product of the numbers of multisets of reads of length $i$ on haplotype $k$ (over all possible values of $i$) gives the number of multisets of reads on haplotype $k$.

**Counting multisets using generating functions.** Suppose we have a multiset $R = \{3 \cdot a, 4 \cdot b, 2 \cdot c\}$ of reads. To count the number of multisets containing exactly six elements from $R$ , where two copies of the same read are indistinguishable, one can use the following method:

1. Write a generating function for each read, e.g. $f_a(x) = 1 + x + x^2 + x^3$.
2. Compute $F(x) = f_a(x) \cdot f_b(x) \cdot f_c(x)$.
3. Extract the coefficient of $x^6$.

The generating function $f_a(x)$ denotes the four available choices concerning read $a$ available when constructing a sub-multiset. We can pick zero, one, two, or three copies of $a$. In the product polynomial $F(x)$, the coefficient of $x^n$ is the number of sub-multisets of $n$ elements.

In our setting, we know that there are $N_{k,i}$ reads of length $i$ on haplotype $k$. There are $G$ possible distinct reads of length $i$ corresponding to the $G$ possible distinct stop positions in haplotype $k$. Some of these reads will be 'valid', i.e. permissible according to conditions defined for $M_{kp}$, where $k$ is the index of the haplotype and $p \in \{1, 2, 3, 4\}$, and others will be 'invalid'. For a fixed choice of haplotype $k$ and length $i$, we write $G$ generating functions $f_{i,j,k}(x)$, for $1 \leq j \leq G$. When estimating the quantities $|M_{11}|, |M_{12}|, |M_{13}|, |M_{14}|$, $|M_{21}|, |M_{22}|, |M_{23}|, |M_{24}|$, some reads may be invalid for some of the quantities. If the read of length $i$ stopping at position $j$ on haplotype $k$ is valid, then the generating function is $f_{i,j,k}(x) = 1 + x + x^2 + \cdots x^{N_{k,i}}$. If the read is invalid, then the generating function is $f_{i,j,k}(x) = x^0$. In the product $F_{k,i}(x) = \prod_{j=1}^{G} f_{i,j,k}(x)$, the coefficient of $x^{N_{k,i}}$, denoted by $a_{N_{k,i}}$, counts the number of multisets of reads of length $i$ from haplotype $k$. The product $\prod_{i=1}^{\lambda_k} a_{N_{k,i}}$ gives the number of multisets of $N_k$ reads arising from haplotype $k$. The roles of haplotypes 1 and 2 can be reversed if $x_1 < x_2$. When $x_1 > x_2$, the conditions for reads in each read sequencing output in $M_{11}, M_{12}, \ldots, M_{23}, M_{24}$ are as follows:

- $M_{11}$:
  1. If $j = x_1$, reads are in group 1(b) if $i \geq x_1$; they are in group 2(b) if $x_1 - x_2 < i < x_1$; they are in group 2(c) if $1 \leq i \leq x_1 - x_2$. All of these are valid reads.
  2. If $x_2 < j < x_1$, reads are in group 1(a) if $i \geq j$; they are in group 2(b) if $j - x_2 < i < j$; they are in group 2(c) if $1 \leq j \leq j - x_2$. All these reads are valid.
  3. If $1 < j \leq x_2$, reads are in group 1(a) if $i \geq j$; they are in group 2(a) if $1 \leq i < j$. All these reads are valid.
  4. If $j = 1$, reads are in group 1(a) for all $i$. All these reads are valid.
  5. If $x_1 < j \leq G$, reads are in group 3(b) if $1 \leq i \leq j - x_2$. All of these are valid reads. All reads of length $i$ with stop position $j$ not mentioned above are invalid.
- $M_{12}$:
  1. If $j = x_1$, reads are in group 1(b) if $i \geq x_1$; they are in group 2(b) if $x_1 - x_2 < i < x_1$; they are in group 2(c) if $1 \leq i \leq x_1 - x_2$. Reads in group 1(b) are invalid. Reads in groups 2(b) and 2(c) are valid.
  2. Same as (2) in $M_{11}$.
  3. Same as (3) in $M_{11}$.
  4. Same as (4) in $M_{11}$.
  5. Same as (5) in $M_{11}$.
- $M_{13}$:
  1. If $j = x_1$, reads are in group 1(b) if $i \geq x_1$; they are in group 2(b) if $x_1 - x_2 < i < x_1$; they are in group 2(c) if $1 \leq i \leq x_1 - x_2$. Reads in groups 1(b) and 2(c) are valid. Reads in group 2(b) are invalid.
  2. If $x_2 < j < x_1$, reads are in group 1(a) if $i \geq j$; they are in group 2(b) if $j - x_2 < i < j$; they are in group 2(c) if $1 \leq j \leq j - x_2$. Reads in groups 1(a) and 2(c) are valid. Reads in group 2(b) are invalid.
  3. Same as (3) in $M_{11}$.
  4. Same as (4) in $M_{11}$.
  5. Same as (5) in $M_{11}$.
- $M_{14}$:

1. If $j = x_1$, reads are in group 1(b) if $i \geq x_1$; they are in group 2(b) if $x_1 - x_2 < i < x_1$; they are in group 2(c) if $1 \leq i \leq x_1 - x_2$. Reads in group 2(c) are valid. Reads in groups 1(b) and 2(b) are invalid.
2. If $x_2 < j < x_1$, reads are in group 1(a) if $i \geq j$; they are in group 2(b) if $j - x_2 < i < j$; they are in group 2(c) if $1 \leq j \leq j - x_2$. Reads in groups 1(a) and 2(c) are valid. Reads in group 2(b) are invalid.
3. Same as (3) in $M_{11}$.
4. Same as (4) in $M_{11}$.
5. Same as (5) in $M_{11}$.

- $M_{21}$:
  1. If $j = x_2$, reads are in group 1(b) if $i \geq x_2$; they are in group 2(a) if $i < x_2$. All of these are valid reads.
  2. If $1 \leq j < x_2$, reads are in group 1(a) if $i \geq j$; they are in group 2(a) if $i < j$. All of these are valid reads.
  3. If $x_2 < j \leq x_1$, reads are in group 2(b) if $j - x_2 < i < j$; they are in group 2(c) if $1 \leq i \leq j - x_2$. All of these are valid reads. If $i \geq j$, these reads would support the heterozygous locus, they are invalid.
  4. If $x_1 < j \leq G$, reads are in group 3(b) if $1 \leq i \leq j - x_2$. All of these are valid reads. All reads of length $i$ with stop position $j$ not mentioned above are invalid.

- $M_{22}$:
  1. If $j = x_2$, reads are in group 1(b) if $i \geq x_2$; they are in group 2(a) if $i < x_2$. Reads in group 2(a) are valid. Reads in group 1(b) are invalid.
  2. Same as (2) in $M_{21}$.
  3. Same as (3) in $M_{21}$.
  4. Same as (4) in $M_{21}$.

- $M_{23}$:
  1. Same as (1) in $M_{21}$.
  2. Same as (2) in $M_{21}$.
  3. If $x_2 < j \leq x_1$, reads are in group 2(b) if $j - x_2 < i < j$; they are in group 2(c) if $1 \leq i \leq j - x_2$. Reads in group 2(c) are valid. Reads in group 2(b) are invalid. If $i \geq j$, these reads would support the heterozygous locus, they are invalid.
  4. Same as (4) in $M_{21}$.

- $M_{24}$:
  1. If $j = x_2$, reads are in group 1(b) if $i \geq x_2$; they are in group 2(a) if $i < x_2$. Reads in group 2(a) are valid. Reads in group 1(b) are invalid.
  2. Same as (2) in $M_{21}$.
  3. If $x_2 < j \leq x_1$, reads are in group 2(b) if $j - x_2 < i < j$; they are in group 2(c) if $1 \leq i \leq j - x_2$. Reads in group 2(c) are valid. Reads in group 2(b) are invalid. If $i \geq j$, these reads would support the heterozygous locus, they are invalid.
  4. Same as (4) in $M_{21}$.

**Minor remark on the calculation of $|S|$.** While counting all valid sequencing outputs $|S|$ in the main text, we use $T$ to denote the number of sequencing outputs having $N_{k,i}$ reads of length $i$ on haplotype $k$ for all $i \in [1, \lambda_k]$ and for all $k \in \{1, 2\}$. $T$ can also be computed directly without using generating functions. Recall that $n$ indistinguishable objects can be placed into $m$ distinguishable bins in $\binom{n+m-1}{n}$ ways. The $N_{k,i}$ number of reads of length $i$ on haplotype $k$ are only distinguishable by their stop positions. Treating the stop positions as distinguishable bins, the number of ways in which the $N_{k,i}$ number of reads can be stored in $G$ distinguishable bins is $\binom{G+N_{k,i}-1}{N_{k,i}}$. This is the number of multisets of reads of length $i$ on haplotype $k$. Thus, the total number of multisets of reads on haplotype $k$ is $\prod_i \binom{G+N_{k,i}-1}{N_{k,i}}$. Accordingly, $T = \prod_{k=1}^{2} \prod_i \binom{G+N_{k,i}-1}{N_{k,i}}$. However, deriving a direct formula for the other quantities, i.e., $T_1, T_2, T_{12}$, $M_{11}, M_{12}, \ldots, M_{24}$, is tedious.

### Supplementary Note S2: Software commands for CGProb

To run CGProb, we use the following commands. Comments are indicated using #.

```
$ python3 scripts/create_genome.py 1000000
# Simulation for the first haplotype
$ seqrequester simulate -truncate -genome myGenome.fasta -genomesize 1000000 -coverage 50 \
-distribution readLengthDistribution.txt > readsHap1.fasta
$ seqrequester summarize -simple readsHap1.fasta > originalReadLengthsHap1.txt
# Simulation for the second haplotype
$ seqrequester simulate -truncate -genome myGenome.fasta -genomesize 1000000 -coverage 50 \
-distribution readLengthDistribution.txt > readsHap2.fasta
$ seqrequester summarize -simple readsHap2.fasta > originalReadLengthsHap2.txt
# Scale down read lengths by factor 1000
$ python3 scripts/scale_down.py originalReadLengthsHap1.txt newShortLengthsHap1.txt 1000
$ python3 scripts/scale_down.py originalReadLengthsHap2.txt newShortLengthsHap2.txt 1000
# Obtain read count on each haplotype
$ tail -2 readsHap1.fasta | head -1 | awk -F'[=,]' '{print $2}' > numReadsHap1.txt
$ tail -2 readsHap2.fasta | head -1 | awk -F'[=,]' '{print $2}' > numReadsHap2.txt
# Computation
$ CGProb/compute -g 1000 -R $(cat numReadsHap1.txt) -r $(cat numReadsHap2.txt) \
-h 200 -D $(cat newShortLengthsHap1.txt) -d $(cat newShortLengthsHap2.txt) -p 128 -t 256 \
> output.txt
$ tail output.txt | grep "probability = " | awk '{print $3}' > probability.txt
```

### Supplementary Note S3: Software commands for RAFT

**Running Hifiasm** We use the following command to run Hifiasm with 32 threads on an input FASTA file.

```
$ hifiasm -o assembly -t 32 input.fasta
```

**Running RAFT-Hifiasm** We use the following sequence of commands to run the RAFT-Hifiasm pipeline with 32 threads on an input FASTA file.

```
$ hifiasm -o errorcorrect -t 32 --write-ec input.fasta 2> errorcorrect.log
$ COVERAGE=$(grep "homozygous" errorcorrect.log | tail -1 | awk '{print $6}')
$ hifiasm -o getOverlaps -t 32 --dbg-ovec errorcorrect.ec.fa 2> getOverlaps.log
$ cat getOverlaps.0.ovlp.paf getOverlaps.1.ovlp.paf > overlaps.paf
$ raft -e ${COVERAGE} -o fragmented errorcorrect.ec.fa overlaps.paf
$ hifiasm -o finalasm -t 32 -r1 fragmented.reads.fasta 2> finalasm.log
```

**Commands for evaluating RAFT using simulated data** We used Seqrequester (commit: `31141c1`) along with a publicly available trio-based assembly of HG002 using Hifiasm to generate simulated read-sequencing datasets. In this example, each haplotype's coverage was set to 25.

```
$ seqrequester simulate -truncate -genome HG002_Trio_Assembly_Hap1.fa -genomesize 2935689000 \
-coverage 25 -distribution read_length_distribution.txt > reads.hap1.fasta
$ seqrequester simulate -truncate -genome HG002_Trio_Assembly_Hap2.fa -genomesize 3033188759 \
-coverage 25 -distribution read_length_distribution.txt > reads.hap2.fasta
$ cat reads.hap1.fasta reads.hap2.fasta > reads.fasta
```

To identify genomic intervals with zero alignment coverage, we use Bedtools (v2.29.1).

```
$ samtools faidx HG002_Trio_Assembly_Hap1.fa
$ samtools faidx HG002_Trio_Assembly_Hap2.fa
$ cut -f1,2 HG002_Trio_Assembly_Hap1.fa.fai > asm.contig.lengths.txt
$ cut -f1,2 HG002_Trio_Assembly_Hap2.fa.fai >> asm.contig.lengths.txt
$ grep ">" reads.fasta > reads.fasta.headers
$ cat reads.fasta.headers | awk -F '[=,-]' '{print $9"\t"$5"\t"$6}' | \
sort -k 1,1 -k2,2n -k3,3nr > reads.bed
$ bedtools genomecov -i reads.bed -g asm.contig.lengths.txt -bga | \
awk '{if ($4==0) print $0}' > zero_cov.bed
```

To identify all the non-contained reads, we first identify the set of contained reads based as reads having 100% alignment identity with a longer read. All-vs-all alignments were computed using minimap2 (v2.26-r1175). The command is as follows.

```
$ minimap2 -t 256 -w 101 -k 27 -g 500 -B 8 -O 8,48 -E 4,2 -cx ava-ont reads.fasta reads.fasta \
> overlaps.paf
$ cat overlaps.paf | awk -v minid=100 '{
    if ($3 == 0 && $2 == $4 && $2 < $7 && $10*100.0/$11 >= minid) print $0
}' | cut -f1 > contained1
$ cat overlaps.paf | awk -v minid=100 '{
    if ($8 == 0 && $7 == $9 && $7 < $2 && $10*100.0/$11 >= minid) print $0
}' | cut -f6 > contained2
$ cat contained1 contained2 | sort | uniq > contained_reads.txt
$ cat reads.fasta.headers | sort | sed 's/>//g' > reads.fasta.sorted.headers
$ comm -23 reads.fasta.sorted.headers contained_reads.txt > non-contained_reads.txt
$ seqtk subseq reads.fasta non-contained_reads.txt > non-contained_reads.fasta
```

We use the set of non-contained reads to identify coverage gaps in the standard string graph formulation as follows.

```
$ minimap2 -t 32 -N 50 -cx map-ont HG002_Trio_Assembly_Hap1.fa non-contained_reads.fasta \
> non-contained_to_asm.paf
$ minimap2 -t 32 -N 50 -cx map-ont HG002_Trio_Assembly_Hap2.fa non-contained_reads.fasta \
>> non-contained_to_asm.paf
$ cat non-contained_to_asm.paf | awk '{
    if ($3 == 0 && $2 == $4 && $2 == $10) print $6"\t"$8"\t"$9
}' | sort -k1,1 -k2,2n -k3,3nr > non-contained.exactmapped.bed
$ bedtools genomecov -i non-contained.exactmapped.bed -g asm.contig.lengths.txt -bga | \
awk '{if ($4==0) print $0}' > asm.uncovered.bed
$ bedtools merge -d 500 -i asm.uncovered.bed > temp.bed && mv temp.bed asm.uncovered.bed
# Identify intervals with coverage gaps due to Contained Read Deletion
$ bedtools subtract -A -a asm.uncovered.bed -b zero_cov.bed > asm.CRD.uncovered.bed
# printEdgeBedIntervals.py identifies intervals of specified length at both ends of contigs
$ python3 printEdgeBedIntervals.py asm.contig.lengths.txt 25000 > genome_25kbp_ends.bed
$ bedtools subtract -A -a asm.CRD.uncovered.bed -b genome_25kbp_ends.bed \
> temp.bed && mv temp.bed asm.CRD.uncovered.bed
$ cat asm.CRD.uncovered.bed | \
awk -F'\t' 'BEGIN{SUM=0}{ SUM+=$3-$2 }END{print SUM}' \
> asm.CRD.uncovered.sum
$ cat asm.CRD.uncovered.bed | awk '{print ($3-$2)}' | \
sort -n > asm.CRD.uncovered.lengths
```

We use the set of non-redundant reads identified by Hifiasm and the alignments of the non-contained reads to identify Hifiasm coverage gaps as follows.

```
$ hifiasm -o assembly -t 32 -r1 reads.fasta
$ grep -P "^A\t" assembly.bp.r_utg.gfa | grep -o -P "read=[^\t]*" | sort \
> assembly.bp.r_utg.gfa.sorted.headers
$ comm -12 assembly.bp.r_utg.gfa.sorted.headers contained_reads.txt \
> assembly.bp.r_utg.gfa.contained.headers
$ seqtk subseq reads.fasta assembly.bp.r_utg.gfa.contained.headers \
> non-redundant.fasta
$ minimap2 -t 32 -N 50 -cx map-ont HG002_Trio_Assembly_Hap1.fa non-redundant.fasta \
> non-redundant.paf
$ minimap2 -t 32 -N 50 -cx map-ont HG002_Trio_Assembly_Hap2.fa non-redundant.fasta \
>> non-redundant.paf
$ cat non-redundant.paf | awk '{
    if ($3 == 0 && $2 == $4 && $2 == $10) print $6"\t"$8"\t"$9
}' > non-redundant.bed
```

```
$ cat non-redundant.paf | awk '{
    if ($8 == 0 && $7 == $9 && $7 == $10) print $6"\t"$8"\t"$9
}' >> non-redundant.bed
$ cat non-redundant.bed | sort -k 1,1 -k2,2n -k3,3nr > non-redundant.exactmapped.bed
$ bedtools subtract -a asm.CRD.uncovered.bed -b non-redundant.exactmapped.bed \
> unresolved_gaps.bed
$ cat unresolved_gaps.bed | \
awk -F'\t' 'BEGIN{SUM=0}{ SUM+=$3-$2 }END{print SUM}' > unresolved_gaps.sum
$ cat unresolved_gaps.bed | awk '{print ($3-$2)}' | sort -n > unresolved_gaps.lengths
```

For the RAFT-Hifiasm method, to identify all the non-contained reads, we first identify the set of contained reads based on fragmented reads having 100% alignment identity with a longer read. The above commands are rerun using fragmented reads instead of the input reads.

**Commands for evaluating RAFT using real data** We used QUAST to estimate **NG50** assuming the size of each haplotype to be 3.1 Gb and the **length of the longest contig**.

```
$ python quast.py -t 256 -o results --est-ref-size 3099922541 --large primary_contigs.fa
$ python quast.py -t 256 -o results --est-ref-size 3099922541 --large assembly_hap1.fa
$ python quast.py -t 256 -o results --est-ref-size 3099922541 --large assembly_hap2.fa
```

**Switch error rate** and **Hamming error rate** were computed using yak trioeval. Complementary parental data from HG003 in the form of Illumina HiSeq data were used to compute k-mer counts in parents. These were compared against k-mers in contigs from phased assemblies.

```
$ zcat HG003_HiSeq30x_subsampled_R1.fastq.gz > HG003.fastq
$ zcat HG003_HiSeq30x_subsampled_R2.fastq.gz >> HG003.fastq
$ zcat HG004_HiSeq30x_subsampled_R1.fastq.gz > HG004.fastq
$ zcat HG004_HiSeq30x_subsampled_R2.fastq.gz >> HG004.fastq
$ yak count -b37 -t 128 -o paternal.yak HG003.fastq
$ yak count -b37 -t 128 -o maternal.yak HG004.fastq
$ yak trioeval -t 128 paternal.yak maternal.yak assembly_hap1.fa > hap1.out
$ yak trioeval -t 128 paternal.yak maternal.yak assembly_hap2.fa > hap2.out
```

To compute **gene completeness statistics**, we used asmgene.

```
$ minimap2 -cxsplice:hq -t 128 grch38_contigs.fa cDNA.fa > grch38_cDNA.paf
$ minimap2 -cxsplice:hq -t 128 assembly_hap1.fa grch38_contigs.fa > hap1_cDNA.paf
$ minimap2 -cxsplice:hq -t 128 assembly_hap2.fa grch38_contigs.fa > hap2_cDNA.paf
$ paftools.js asmgene -a grch38_cDNA.paf hap1_cDNA.paf > hap1.stats
$ paftools.js asmgene -a grch38_cDNA.paf hap2_cDNA.paf > hap2.stats
```

**Telomere-to-Telomere contigs** were computed using the HPRC workflow at `https://github.com/biomonika/HPP/blob/main/assembly/wdl/workflows/assessAsemblyCompletness.wdl`. CHM13v2 was used as the reference genome for the workflow. The resulting T2T contigs summarised by the workflow were vetted to ensure that the contigs were within 10% of the length of the corresponding chromosome in CHM13v2.

**QV** was obtained using yak. Complementary HiC data for HG002 was used to compute k-mer statistics as a comparison against the assembly contigs. Both sets of phased assembly contigs were passed as input for the comparison.

```
$ yak count -b37 -y 128 -o complementary-kmers.yak hg002_hic.fq
$ yak qv -t 128 -p -K3.2g -l100k complementary-kmers.yak assembly_hap1hap2.fa
```

**Variant calling statistics**, i.e. SNPs and INDELs were first called using dipcall and the VCF file was compared against the GIAB variant calling benchmark v4.2.1 for HG002 (treated as ground truth).

```
$ dipcall.kit/run-dipcall -x dipcall.kit/hs38.PAR.bed output \
GRCh38_no_alt_analysis_set.fna assembly_hap1.fa assembly_hap2.fa > output.mak
$ make -j4 -f output.mak
$ tabix -p vcf output.dip.vcf.gz
$ bedtools intersect -a output.dip.bed -b HG002_GRCh38_v4.2.1_noinconsistent.bed \
> conf_giab_restrict.bed
$ hap.py HG002_GIAB_v4.2.1.vcf output.dip.vcf.gz -f conf_giab_restrict.bed \
-r GRCh38_no_alt_analysis_set.fna -o compare --engine=vcfeval --pass-only
```

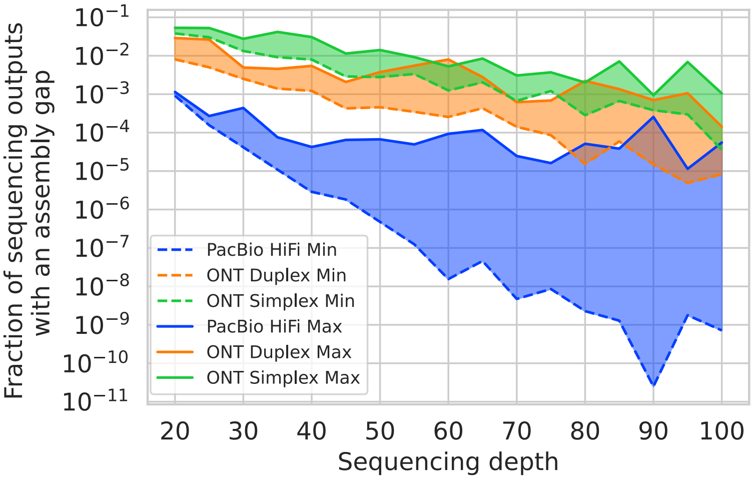


**Figure S1: Frequency of observing an assembly gap near a germline heterozygous locus**. In the main text Figure 1B, we showed the median frequencies of observing an assembly gap near a germline heterozygous variant locus in a string graph. For each choice of sequencing technology and sequencing depths, the median value was reported by running CGProb using five simulated read length distributions. From the same experiment, the above figure shows the minimum and maximum values.

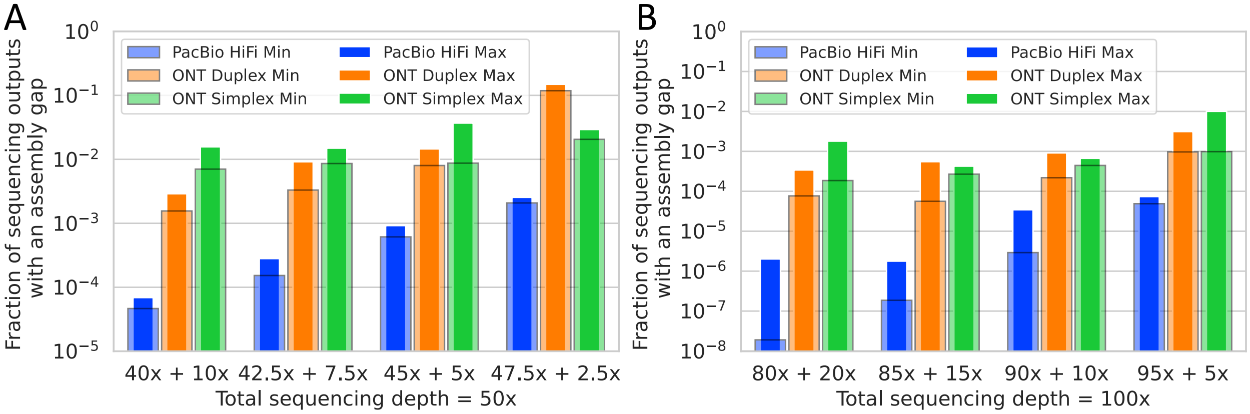


**Figure S2: Frequency of observing an assembly gap near a somatic heterozygous locus**. In the main text Figures 1C and 1D, we showed the median frequencies of observing an assembly gap near a somatic heterozygous variant locus in a string graph. For each choice of sequencing technology and sequencing depths, the median value was reported by running CGProb using five simulated read length distributions. From the same experiment, the above figures show the minimum and maximum values.

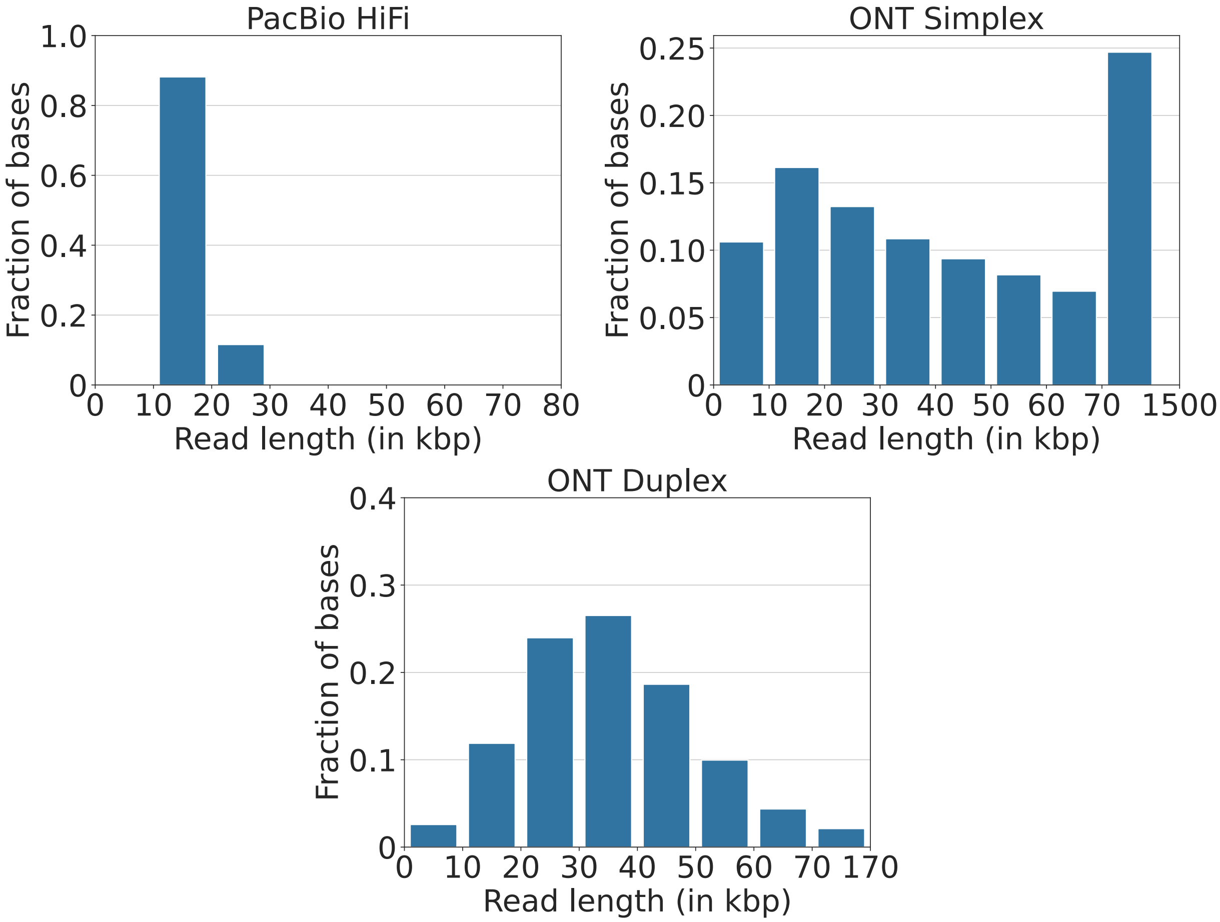


**Figure S3: Read length distributions corresponding to the different read sequencing technologies.** PacBio HiFi dataset has reasonably uniform read lengths, whereas the range of read lengths in ONT Simplex and ONT Duplex datasets is large.

| | Dataset | State | Read Count | Bases (Gb) | N50 (kb) | Longest read (kb) |
|---|---|---|---|---|---|---|
| Simulated datasets | HiFi (30x) | Original | 4.2M | 89.5 | 21.3 | 48.7 |
| | | Fragmented | 6.7M | 90.8 | 20.0 | 47.4 |
| | ONT Simplex (30×) | Original | 5.4M | 89.5 | 39.6 | 572.4 |
| | | Fragmented | 7.9M | 90.8 | 20.0 | 175.1 |
| | ONT Simplex (50×) | Original | 8.9M | 149.2 | 39.7 | 572.4 |
| | | Fragmented | 13.2M | 151.4 | 20.0 | 189.5 |
| | ONT Duplex (30×) | Original | 3.4M | 89.5 | 34.1 | 170.6 |
| | | Fragmented | 6.1M | 90.9 | 20.0 | 120.7 |
| | ONT Duplex (50×) | Original | 5.7M | 149.2 | 34.1 | 170.6 |
| | | Fragmented | 10.2M | 151.5 | 20.0 | 118.9 |
| Publicly available real sequencing datasets | D1: HiFi (36×) | Original | 7.4M | 110.5 | 14.7 | 46.9 |
| | | Fragmented | 7.9M | 110.8 | 14.6 | 38.0 |
| | D2: ONT Duplex (32×) | Original | 3.7M | 99.3 | 34.2 | 170.6 |
| | | Fragmented | 6.8M | 100.9 | 20.0 | 95.9 |
| | D3: HiFi (36×) + ONT Duplex (32×) | Original | 11.1M | 209.9 | 19.3 | 170.6 |
| | | Fragmented | 14.7M | 211.7 | 17.8 | 95.9 |
| | D4: ONT high-acc UL (40×) | Original | 5.6M | 126.9 | 91.7 | 1397.8 |
| | | Fragmented | 10.3M | 129.3 | 20.5 | 397.1 |

**Table S1: Read length statistics of sequencing datasets used for benchmarking RAFT algorithm.** The first five read datasets were simulated from a long-read Trio-based assembly of the HG002 human sample. D1, D2, and D4 are publicly available real datasets. D3 is constructed by merging D1 and D2. We used RAFT on the original datasets to produce the fragmented datasets. Read length statistics are shown for both the original and their fragmented versions.

| Dataset | Method | SNPs | | | INDELs | | | Longest | QV | Hamming |
|---|---|---|---|---|---|---|---|---|---|---|
| | | Recall (%) | Precision (%) | F1 (%) | Recall (%) | Precision (%) | F1 (%) | Contig (Mbp) | | Error (%) |
| D1: HiFi (36×) | Hifiasm | 99.04 | 99.81 | 99.43 | 97.49 | 97.82 | 97.65 | 176.5/138.5 | 52.2/52.4 | 26.0/23.9 |
| | RAFT-Hifiasm | 99.05 | 99.86 | 99.46 | 97.45 | 97.86 | 97.65 | 140.3/133.9 | 52.4/51.1 | 24.7/24.5 |
| D2: Duplex (ONT) (32×) | Hifiasm | 98.36 | 99.63 | 98.99 | 86.02 | 69.91 | 77.14 | 180.0/137.6 | 47.7/47.9 | 26.7/17.9 |
| | RAFT-Hifiasm | 99.10 | 99.68 | 99.39 | 86.48 | 70.35 | 77.58 | 201.1/174.7 | 48.0/48.4 | 24.3/22.1 |
| D3: HiFi (36×) + Duplex (ONT) (32×) | Hifiasm | 97.71 | 99.76 | 98.72 | 95.58 | 96.19 | 95.89 | 137.7/148.5 | 51.2/50.5 | 19.2/23.6 |
| | RAFT-Hifiasm | 99.21 | 99.79 | 99.50 | 96.96 | 96.38 | 96.67 | 140.3/192.5 | 50.9/52.4 | 23.9/26.2 |
| D4: High-acc UL (ONT) (40×) | Hifiasm | 57.87 | 97.66 | 72.67 | 51.04 | 67.57 | 58.16 | 77.2/103.6 | 40.9/41.1 | 8.1/5.8 |
| | RAFT-Hifiasm | 98.36 | 99.51 | 98.93 | 87.92 | 72.82 | 79.66 | 140.9/144.9 | 47.1/47.1 | 26.7/22.1 |

**Table S2: Additional quality statistics of the HG002 phased assemblies produced by RAFT-Hifiasm and Hifiasm.** We measured the base-level accuracy of the assembled sequences by using SNP and indel calls available from Genome in a Bottle benchmark (v4.2.1). We used Dipcall to call variants from the assembled sequences. We restricted our evaluation to the GIAB's high-confidence intervals. Information about the longest contig was extracted using QUAST. QV and Hamming error were estimated using yak. These genome assemblies should be interpreted as partially phased assemblies [16] because we did not use parental or Hi-C data during assembly.

| **Software** | **Version** | **Purpose** |
|---|---|---|
| Bedtools | 2.29.1 | Identifying genomic intervals without coverage |
| CGProb | Commit `0f16cfc` | Fraction of sequencing outputs with an assembly gap |
| Dipcall | 0.3 | Variant calling |
| Hap.py | 0.3.9 | Comparing VCF files |
| Minimap2 | 2.26-r1175 | Read alignment, Contig alignment |
| Hifiasm | 0.19.8-r603 | Genome assembly, RAFT-Hifiasm pipeline |
| QUAST | 5.2.0 | NG50 |
| RAFT | Commit `f1e34ad` | Fragmenting long reads |
| Samtools | 1.6 | Indexing reference genomes |
| Seqrequester | r90 (commit ID: 31141c1) | Generating simulated reads |
| Seqtk | 1.3-r106 | Selecting subset of FASTA files |
| Tabix | 1.9 | Indexing VCF files |
| Yak | 0.1-r69-dirty | Switch error, Hamming error, QV |

**Table S3: Software versions.**
